# Local signals in mouse horizontal cell dendrites

**DOI:** 10.1101/143909

**Authors:** Camille A. Chapot, Christian Behrens, Luke E. Rogerson, Tom Baden, Sinziana Pop, Philipp Berens, Thomas Euler, Timm Schubert

## Abstract

The mouse retina contains a single type of horizontal cell, a GABAergic interneuron that samples from all cone photoreceptors within reach and modulates their glutamatergic output via parallel feedback mechanisms. Because horizontal cells form an electrically-coupled network, they have been implicated in global signal processing, such as large scale contrast enhancement. Recently, it has been proposed that horizontal cells can also act locally at the level of individual cone photoreceptors. To test this possibility physiologically, we used two-photon microscopy to record light stimulus-evoked Ca^2+^ signals in cone axon terminals and horizontal cell dendrites as well as glutamate release in the outer plexiform layer. By selectively stimulating the two mouse cone opsins with green and UV light, we assessed whether signals from individual cones remain “isolated” within horizontal cell dendritic tips, or whether they spread across the dendritic arbour. Consistent with the mouse‘s opsin expression gradient, we found that the Ca^2+^ signals recorded from dendrites of dorsal horizontal cells were dominated by M- and those of ventral horizontal cells by S-opsin activation. The signals measured in neighbouring horizontal cell dendritic tips varied markedly in their chromatic preference, arguing against global processing. Rather, our experimental data and results from biophysically realistic modelling support the idea that horizontal cells can process cone input locally, extending the “classical” view of horizontal cells function. Pharmacologically removing horizontal cells from the circuitry reduced the sensitivity of the cone signal to low frequencies, suggesting that local horizontal cell feedback shapes the temporal properties of cone output.

**Highlights:**

- Light-evoked Ca^2+^ signals in horizontal cell dendrites reflect opsin gradient
- Chromatic preferences in neighbouring dendritic tips vary markedly
- Mouse horizontal cells process cone photoreceptor input locally
- Local horizontal cell feedback shapes the temporal properties of cone output

**eTOC Blurb:** Chapot et al. show that local light responses in mouse horizontal cell dendrites inherit properties, including chromatic preference, from the presynaptic cone photoreceptor, suggesting that their dendrites can provide “private” feedback to cones, for instance, to shape the temporal filtering properties of the cone synapse.

## Introduction

Most neurons in the brain have elaborate dendritic arbours capable of more than simply integrating synaptic input. Studies of neurons from different brain regions, such as cerebellar Purkinje cells [1], cortical pyramidal cells [2], hippocampal neurons [3], and retinal amacrine cells [4,5], have demonstrated that dendrites can be functionally highly compartmentalized. Often, multiple dendritic units can both process synaptic input and generate synaptic output independently and at a local scale (reviewed in [6]). The cellular mechanisms supporting dendritic processing include anatomical specialisations, differential distribution of active channels, and the local restriction of intracellular signalling (reviewed in [6]). Moreover, computational work suggests that dendrites can even switch between local and global signal processing, e.g. depending on the stimulus strength [7]. Such functional compartmentalisation of dendritic arbours greatly increases the computational power of single neurons and, therefore, that of the brain.

In the retina, dendritic processing has been mainly studied in ganglion cells [8,9] and amacrine cells [4], where dendritic subunits vary dramatically in size and function: For example, starburst amacrine cell dendritic arbours are divided in sections that individually compute direction of visual motion [10,11], while individual dendritic varicosities of A17 amacrine cells provide local reciprocal feedback to individual rod bipolar cell terminals under low-light conditions [4]. However, also the outer retina contains a candidate for dendritic processing, the horizontal cell (HC). This is a GABAergic interneuron that provides reciprocal feedback to photoreceptors and shapes their transmitter release [12–14]. The dendrites of HCs contact cone photoreceptors (cones), whereas their axon terminal system - when they feature one - contacts rod photoreceptors (rods) [15].

Traditionally, HCs have been implicated in global processing, such as contrast enhancement and the generation of antagonistic centre-surround receptive fields (reviewed in [16]). This is consistent with the fact that HCs form a gap junction-coupled network [17], which allows averaging signals across many cones. However, recent studies suggest that HCs support also a local “mode of operation” and that HC feedback can act at the level of a single synaptic contact between a HC dendritic tip and a cone ([14,18]; for discussion see [19]) (Fig. 1A,B).

**Figure 1.**
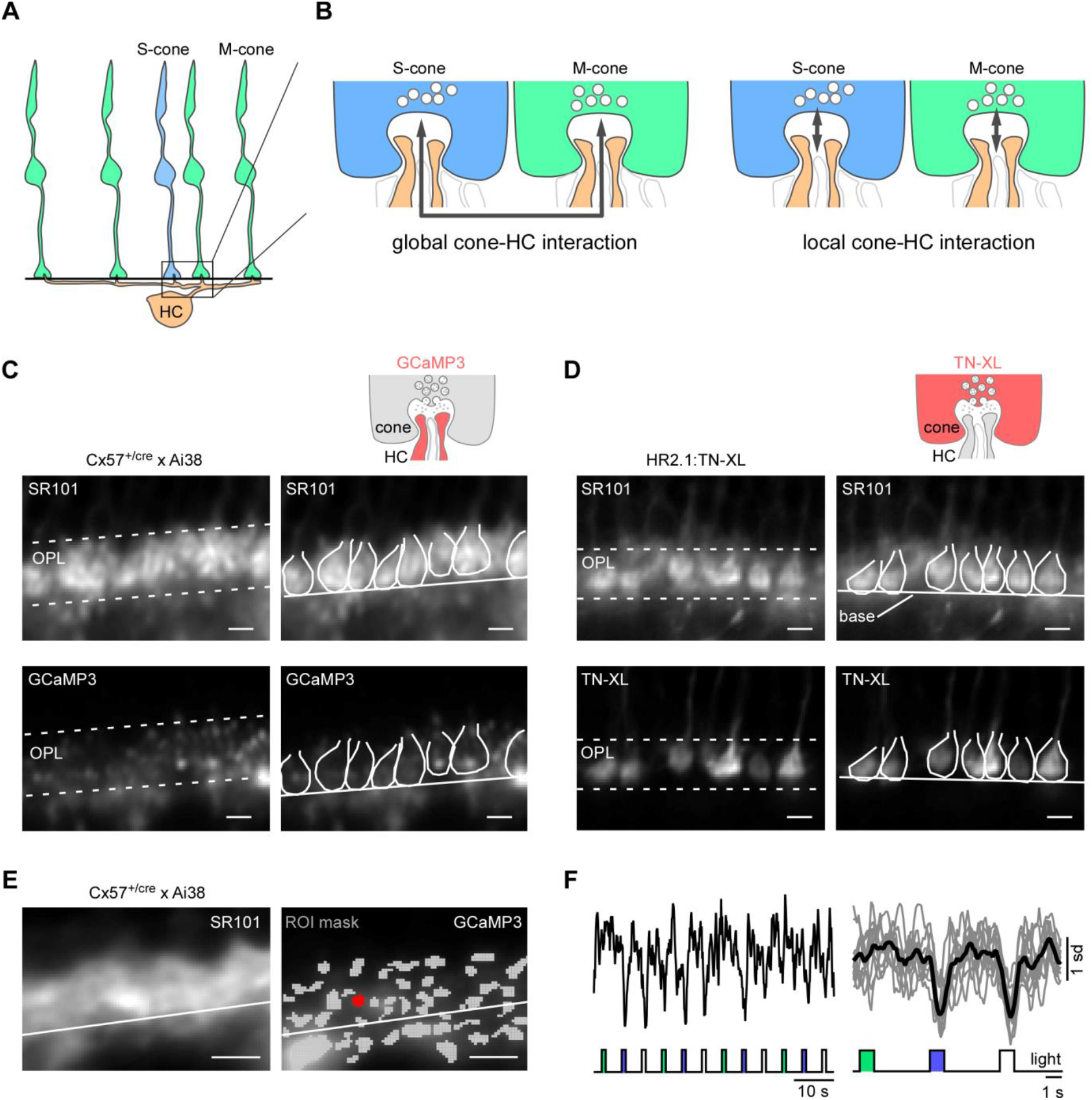
Identification of cone axon terminals and HC processes in mouse retinal slices. **A.** Schematic representation of the connectivity between S-(blue) or M-cones (green) and a horizontal cell (HC, orange). The box corresponds to the enlarged schemata shown in **B.** The black line indicates the cone axon terminal base (as shown in C-E). B. Neighbouring S-(blue) and M-cones (green) with postsynaptic HC dendrites (orange). Bipolar cell dendrites are shown in white. The arrows indicate the hypothesized spread of signals in HC dendrites. *Left:* Global (lateral) signal spread along HC dendrites. *Right:* Local signal processing in HC dendritic tips. **C,D.** Bath application of sulforhodamine 101 (SR101) (top images in C,D) to identify cone axon terminals in retinal slices of the Cx57^+/cre^ x Ai38 (C) and HR2.1:TN-XL (D) mouse lines. Outlines of cone axon terminals were manually drawn for illustration purposes; solid lines indicate cone axon terminal base; dotted lines indicate outer plexiform layer (OPL) borders. Upper right diagram depicts imaged synaptic compartment and biosensor used (red). **E.** *left:* SR101 fluorescence with line marking the cone axon terminal base (analogous to A, C and D). *Right*: GCaMP3-labeled HC processes superimposed by regions of interest (ROIs; grey, exemplary ROI marked red) automatically determined (Methods). **F.** Ca^2+^ responses to green, UV and “white” (GUW) 1-s light flashes of exemplary ROI (in E); continuous Ca^2+^ trace (left) and average of n=10 trials for each stimulus condition (right) are shown (Ca^2+^ signals de-trended by high-pass filtering at ~0.1 Hz and z-normalized, Methods). Scale bars, 5 µm.

Here, we test this idea by recording light stimulus-evoked signals at the HC-cone synapse in a slice preparation of the mouse retina using two-photon Ca^2+^ [20,21] and glutamate imaging [22]. We exploited the particular retinal distribution of mouse cone types to discriminate between global and local processing: Mice express two types of cone opsins, a short (S, “UV") and a medium (M, "green") wavelength-sensitive opsin. So-called "true” S-cones [23] exclusively express S-opsin and are homogenously distributed across the retina, while M-cones co-express both opsins at a ratio that changes from M- to S-opsin-dominant along the dorso-ventral axis [24]. Thus, recording at different retinal locations with different-wavelength stimuli makes it possible to test if signals of neighbouring cones “mix” in the postsynaptic HC dendritic process. We found that cone signals indeed remain local in the contacting HC dendritic tips, suggesting that HCs support a local mode of operation.

## Results

### Identification of cone axon terminals and horizontal cell processes in the mouse retinal slice

We recorded Ca^2+^ signals in retinal slices prepared from transgenic mice (Cx57^+/cre^ x Ai38; see Discussion), which express the Ca^2+^ biosensor GCaMP3 in HCs under the control of the promoter for the gap junction-forming connexin57 (Cx57). HC processes could be identified in retinal slices by their GCaMP3 expression (Fig. 1C). To identify cone axon terminals, we bath-applied SR101 [25], which is taken up from the extracellular solution by the terminals of synaptically very active cells, such as photoreceptors, during vesicle endocytosis [26]. We confirmed that the (larger) SR101-labelled structures in the outer plexiform layer (OPL) were cone axon terminals with slices prepared from HR2.1:TN-XL mice [21], in which exclusively cones express TN-XL (Fig. 1D).

### Light-evoked Ca^2+^ signals in horizontal cell processes

To record light-evoked Ca^2+^ signals in HC dendritic segments, we imaged fields in the OPL while presenting green, UV or “white” light flashes ("GUW protocol", Methods) (Fig. 1E,F). The resulting GCaMP3 fluorescence image series was averaged to anatomically defined regions-of-interest (ROIs) (Methods). We only considered ROIs that responded to white flashes and fulfilled two strict quality criteria, a quality index *(Qi)* and a consistency index *(Ci)* (Suppl. Fig. 1; Methods), yielding 423 ROIs (4.3% from a total of 9,912 ROIs) with reliable light-evoked Ca^2+^ signals for further analysis (Suppl. Fig. 1A-C).

Because the structural layout of the cone synapse is highly stereotypical [27], we assumed that ROIs located close to the cone axon terminal base are likely to be HC dendritic tips, since this is where they make invaginating contacts with the cones (reviewed in [19]). ROIs well above the cone base are expected to belong mostly to HC axon terminal tips (contacting rods), whereas ROIs below the cone base should be located on HC primary dendrites [27]. To get an estimate of each ROI’s identity, we manually determined the base of the cone terminals as a “landmark” (solid lines in Fig. 1A,C-E) in each imaged field based on SR101 labelling. We used the sharp transition between the brightly stained cone axon terminals and the dimmer SR101-labeling below, which likely represents HC dendrites [26]. We estimated the distance *(d_base_)* to the cone axon terminal base for each ROI. Responsive ROIs were most frequent just above the cone axon terminal base (61.5% ROIs within 0 < *d_base_*< 5 µm), within the OPL band occupied by cone terminals. Here, ROIs had the highest *Qi* values (Suppl. Fig. 1D) and the largest light-evoked Ca^2+^ signals (Suppl. Fig. 1E), suggesting that we can indeed measure responses from HC dendritic processes in or very close to the cone synapse.

### Mechanisms underlying light-evoked Ca^2+^ responses in HCs

To confirm that the Ca^2+^ responses were mediated by glutamate release from photoreceptors, we puff-applied the AMPA/KA-type glutamate receptor antagonist NBQX (200 µM) while presenting white flashes (Fig. 2A,B). NBQX significantly decreased the Ca^2+^ baseline level (F0) in HC processes and virtually abolished light-evoked Ca^2+^ signals, as indicated by a significant reduction in response amplitude (*ΔF*) and area-under-the-curve (*F_Area_*) (Fig. 2C-E, for statistics, see Table 1).

**Figure 2.**
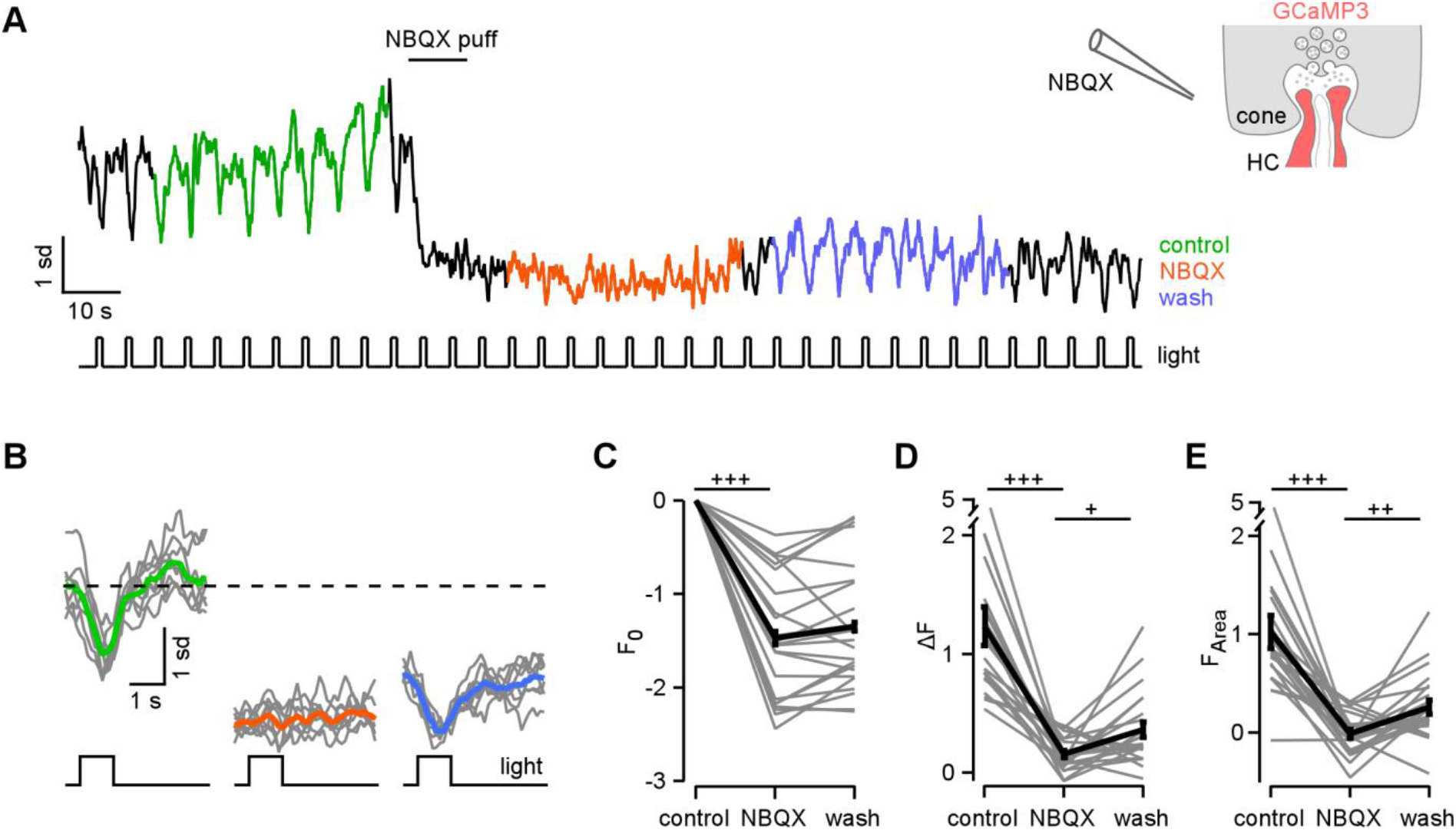
Light-evoked Ca^2+^ responses in HC processes are mediated by activation of AMPA/kainate-type glutamate receptors. **A.** Exemplary Ca^2+^ response of a HC process to white flashes before (control), after a NBQX puff and during wash-out. **B.** Averaged responses for control (green), NBQX (orange) and wash (blue) (trials in grey). **C-E.** Quantification of NBQX effects on response baseline (F_0_, C), amplitude (*ΔF, D*), and area-under-the-curve (*F_Area_*, E) (average of n=23 ROIs from 4 slices, 2 animals). Error bars indicate SEM. +, p < 0.025; ++, p < 0.005; +++, p < 0.0005 (Bonferroni corrected significance threshold).

**Table 1.**
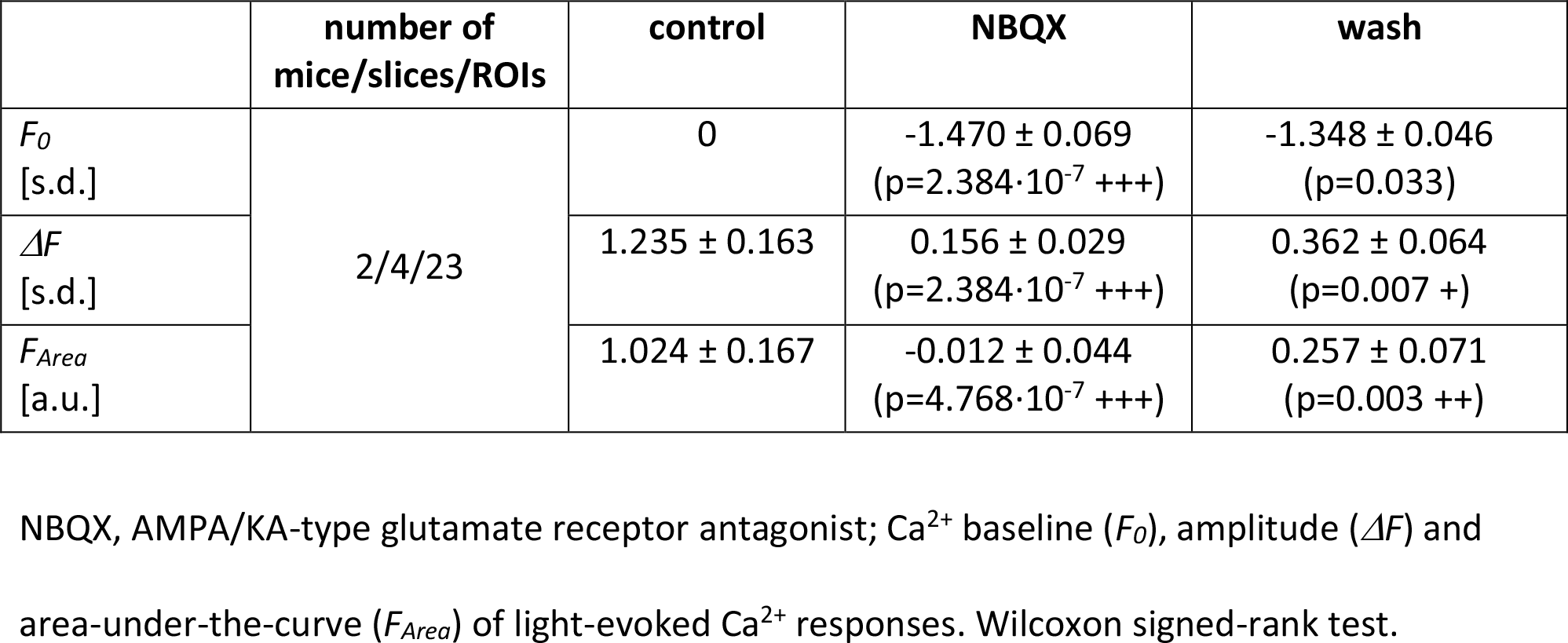
Pharmacology for AMPA/KA-type glutamate receptors

Earlier experiments on isolated mouse HCs had shown that intracellular Ca^2+^ is modulated by influx through Ca^2+^ permeable AMPA/KA receptors, L-type voltage-gated Ca^2+^ channels (VGCCs) and by release from internal Ca^2+^ stores [28]. To test how these pathways contributed to the Ca^2+^ signals in HC dendrites, we puff-applied a mixture of AMPA (50 µM) and KA (25 µM) before and in the presence of blockers (Suppl. Fig. 2). The response amplitudes to AMPA/KA puffs alone decreased during the experiment (Suppl. Fig. 2A,C), possibly caused by downregulation of VGCCs and/or Ca^2+^ stores due to the strong pharmacological stimulus. We estimated this rundown from two consecutive puffs by calculating the ratio of the response amplitudes (*ΔF*_2_*/ΔF*_1_). When applying the L-type VGCC blocker verapamil (100 µM) 5 min before the second AMPA/KA puff, *ΔF_2_/ΔF_1_* was significantly reduced compared to control (Suppl. Fig. 2A,B,E; for statistics, see Table 2), confirming that VGCCs contributed to the signals.

**Table 2.**
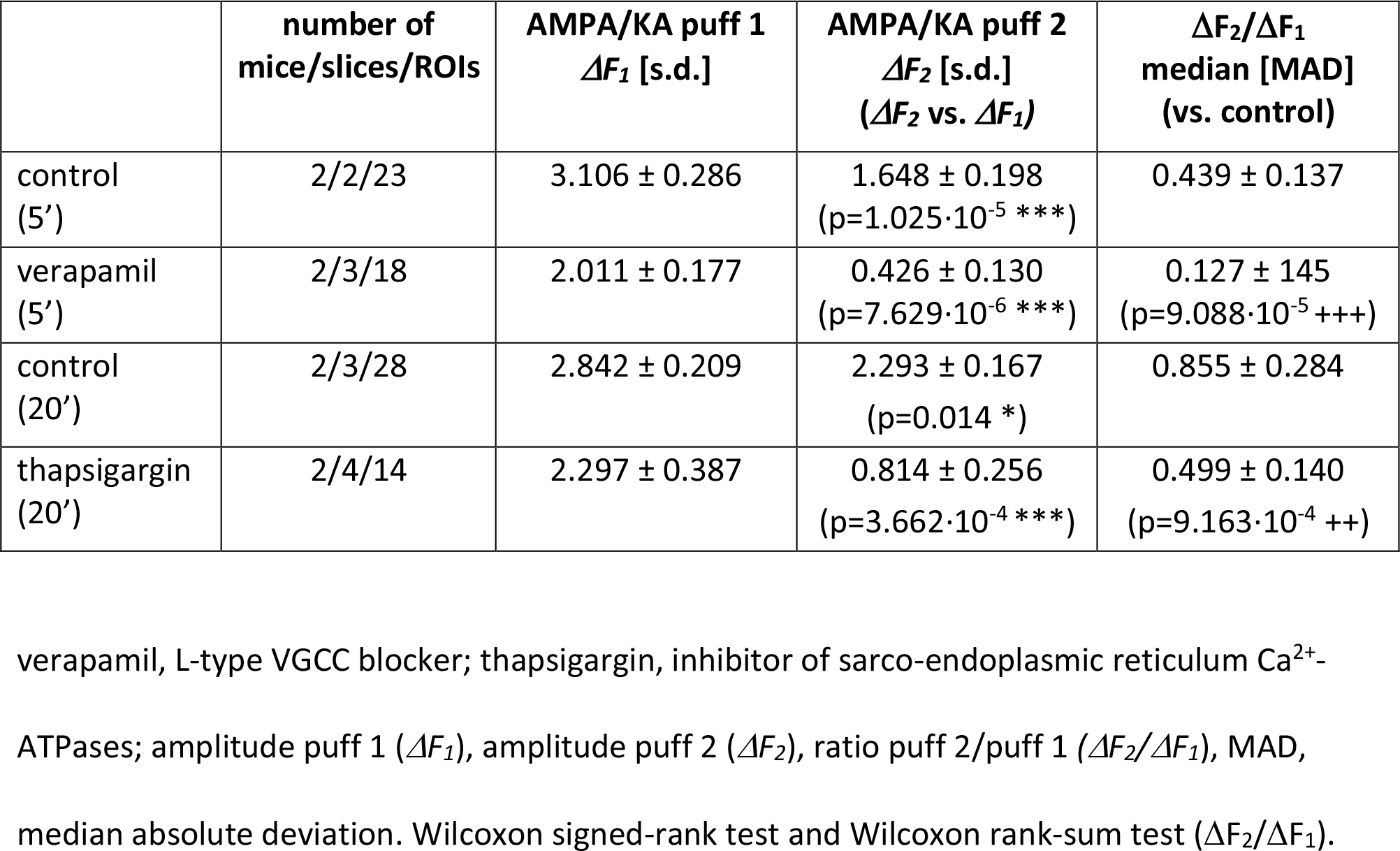
Pharmacology to block voltage-gated Ca^2+^ channels and Ca^2+^ release from stores

| | number of mice/slices/ROIs | AMPA/KA puff 1<br>$\Delta F_1$ [s.d.] | AMPA/KA puff 2<br>$\Delta F_2$ [s.d.]<br>($\Delta F_2$ vs. $\Delta F_1$) | $\Delta F_2/\Delta F_1$<br>median [MAD]<br>(vs. control) |
| --- | --- | --- | --- | --- |
| control<br>(5') | 2/2/23 | $3.106 \pm 0.286$ | $1.648 \pm 0.198$<br>($p=1.025 \cdot 10^{-5}$ ***) | $0.439 \pm 0.137$ |
| verapamil<br>(5') | 2/3/18 | $2.011 \pm 0.177$ | $0.426 \pm 0.130$<br>($p=7.629 \cdot 10^{-6}$ ***) | $0.127 \pm 0.145$<br>($p=9.088 \cdot 10^{-5}$ +++) |
| control<br>(20') | 2/3/28 | $2.842 \pm 0.209$ | $2.293 \pm 0.167$<br>($p=0.014$ *) | $0.855 \pm 0.284$ |
| thapsigargin<br>(20') | 2/4/14 | $2.297 \pm 0.387$ | $0.814 \pm 0.256$<br>($p=3.662 \cdot 10^{-4}$ ***) | $0.499 \pm 0.140$<br>($p=9.163 \cdot 10^{-4}$ ++) |

verapamil, L-type VGCC blocker; thapsigargin, inhibitor of sarco-endoplasmic reticulum $Ca^{2+}$ -ATPases; amplitude puff 1 ( $\Delta F_1$ ), amplitude puff 2 ( $\Delta F_2$ ), ratio puff 2/puff 1 ( $\Delta F_2/\Delta F_1$ ), MAD, median absolute deviation. Wilcoxon signed-rank test and Wilcoxon rank-sum test ( $\Delta F_2/\Delta F_1$ ).

In separate experiments, we tested if intracellular Ca^2+^ stores could be involved in amplifying Ca^2+^ signals in HC processes. We bath-applied the sarco-endoplasmic reticulum Ca^2+^ ATPase (SERCA) inhibitor thapsigargin (5 µM), which blocks Ca^2+^ store refill and leads to depletion of Ca^2+^ stores [28], 20 min before the second AMPA/KA puff. Thapsigargin decreased *ΔF*_2_*/ΔF*_1_ significantly (Suppl. Fig. 2C,D,F, Table 2), suggesting that release from stores contributes to Ca^2+^ signals in HC dendrites.

In summary, the observed light-evoked Ca^2+^ signals in HC processes result from a combination of Ca^2+^ sources, including Ca^2+^-permeable glutamate receptors, VGCCs and release from Ca^2+^ stores, and are modulated by GABA auto-reception (for details, see Suppl. Material, Suppl. Fig. 3 and Suppl. Table 1), in agreement with earlier findings [13,28-31].

**Figure 3.**
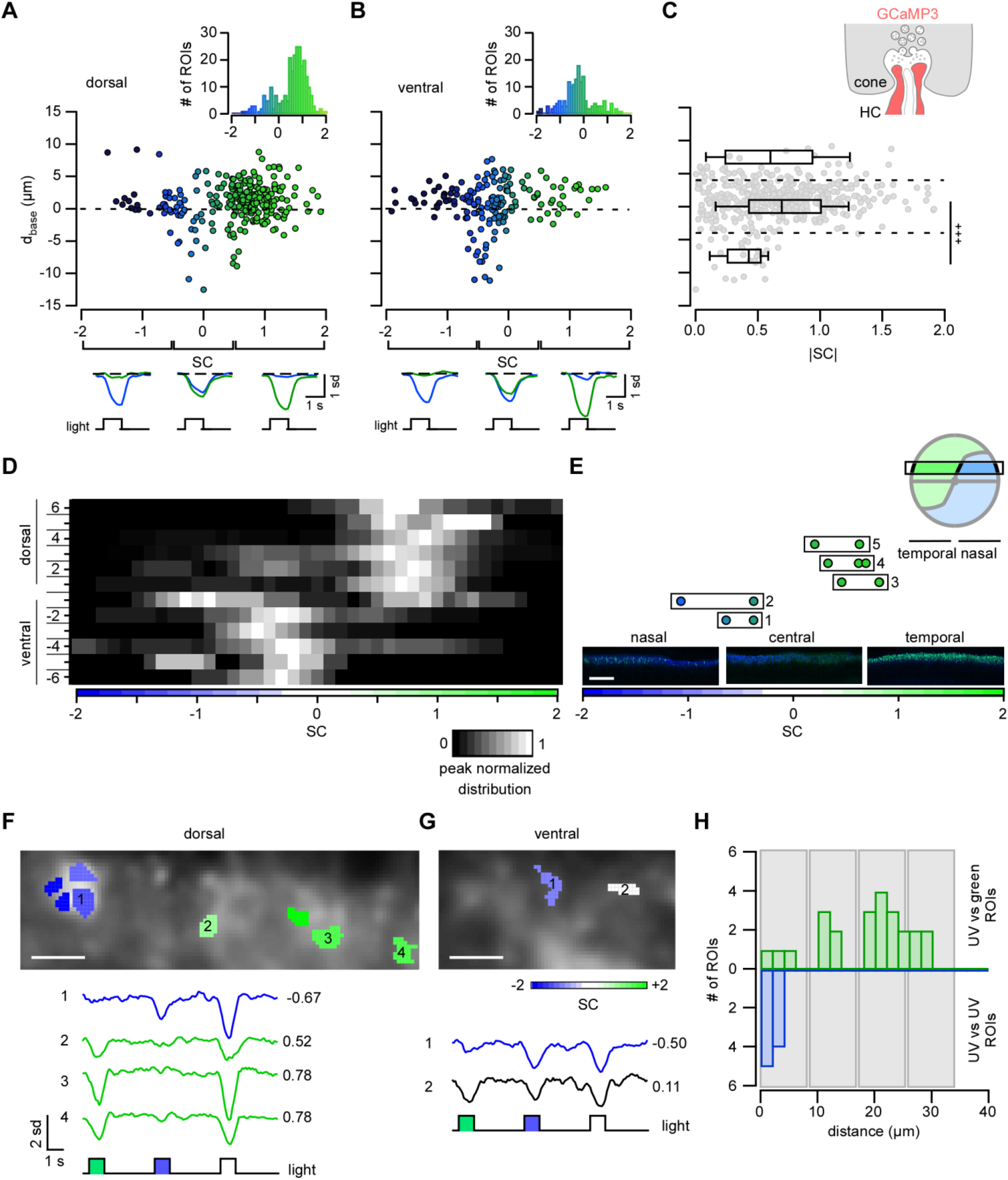
Light-evoked Ca^2+^ signals in HC dendrites reflect the dorso-ventral cone opsin expression gradient and local cone input. **A,B.** Plots showing distance between ROI and cone axon terminal base (*d_base_*) as a function of spectral contrast (SC, Methods) for dorsal (n=262 ROIs) (A) and ventral retina (n=161) (B). Insets: histograms of *SC* distributions. Below: Averaged Ca^2+^ signals in response to green and UV flashes for different *SC* intervals (averages of n=10 trials). **C.** ROI distance to cone axon terminal base (*d_base_*) as a function of |SC| (ROIs from dorsal and ventral retina). ROIs were separated into three groups (dashed lines) depending on *d_base_:* above (*d_base_* > 4 µm), below (*d_base_*< -4 µm), and near the cone axon terminal base (-4 < *d_base_* < 4 µm). **D.** *SC* distribution sorted by retinal slice position (dorsal to ventral; distributions peak normalized for each slice position). **E.** *SC* of ROIs from 5 locations on the same slice (boxes 1-5) along the naso-temporal axis (position +3, see D) and corresponding S-(blue) and M-opsin (green) immunolabeling in the temporal, central and nasal region. **F,G.** Examples of recording fields containing ROIs with different *SC* for dorsal (F) and ventral (G) retina; respective Ca^2+^ signals below (averages of n=10 trials). Colours reflect *SC* preference of each ROI (colour bar in G). **H.** Spatial distribution of UV-(top) and green-(bottom) preferring ROIs relative to each UV ROI (at 0 µm) (for ROIs with |SC| > 0.3; n=22 ROIs from 7 fields, 4 dorsal and 3 ventral retinas). Grey boxes illustrate expected location of neighbouring cone axon terminals. +++, p ≤ 0.0005 (Bonferroni corrected significance threshold). Scale bars, 200 µm in E, 5 µm in F, G.

### Light-evoked Ca^2+^ signals in HCs reflect the dorsal-ventral opsin expression gradient

Next we recorded HC light-evoked Ca^2+^ responses at different locations along the dorso-ventral axis of the retina, using the mouse’ opsin expression gradient as a “tool” to specifically activate different combinations of S-and M-cones. While the mouse retina contains only ~5% “true” Scones [23], ontogenetic M-cones in the ventral retina co-express large amounts of S-opsin and, thus, are “functional” S-cones [24,32]. Therefore, if the spectral preference of the HC Ca^2+^ signals reflects this gradient, this indicates that cones (and not rods) dominantly drive these signals and that we are recording from HC dendrites.

We determined the spectral contrast (SC, Methods) of each ROI as a function of its location along the dorso-ventral axis (Fig. 3). Consistent with the reported opsin gradient [32], we found that dorsal HC responses were dominated by M- and ventral HC responses by S-opsin activation (Fig. 3A,B). ROIs located close to the cone axon terminal base (-4 ≤ *d_base_* ≤ 4 µm) had significantly higher absolute SC values (|*SC_-4…+4_*|=0.717 ± 0.022, n=342) than ROIs below (*d_base_* < -4 µm, |*SC*_<*4*_|=0.417 ± 0.045, n=28, p=1.611·10^-5^, Wilcoxon rank-sum test) (Fig. 3C). This suggests that the HC distal tips reflect the contacted cone’s chromatic preference and, thus, local signals. More proximal dendrites, on the other hand, average across cones, and thus, show spatial integration, in agreement with the “funnel” shape of the *d_base_* vs. *SC* plot (Fig. 3A,B). In the transitional zone between dorsal and ventral retina halves, both a UV- and a green-dominated ROI population co-exist (Fig. 3D). Opsin immunostaining of slices from this zone confirmed that the distribution of UV and green ROIs reflects cone opsin expression (Fig. 3E): ROIs in the nasal part of the slice were UV-sensitive, those in the temporal part were green-sensitive, consistent with the transitional zone running at a shallow angle relative to the naso-temporal axis (Fig. 3E, right scheme) [32]. Together, our data indicate that the activity recorded in ROIs close to the cone axon terminal base is mostly cone-driven and likely reflects activity in HC dendritic tips.

### Local light-evoked Ca^2+^ signals in HC dendritic tips

Next we assessed if signals from individual cones remain “isolated” within HC distal dendrites or if they spread across the cells' dendritic arbours (or the electrically coupled HC network) (Fig. 1B). We looked for recording fields where neighbouring ROIs have different *SC* preferences (i.e. contain ROIs with *SC* > 0 and ROIs with *SC* < 0). Indeed, this was the case for 15 out of a total of 125 recording fields in both dorsal (5 fields; Fig. 3F) and ventral retina (10 fields; Fig. 3G).

To quantify this finding, we focused on “purely” UV and green ROIs (|SC| > 0.3; 7 fields, 43 ROIs) and analysed the distribution of the lateral distance between each UV ROI and its neighbours (Fig. 3H). We found that UV ROIs clustered in close proximity (< 10 µm) of each UV ROI – suggesting that they are driven by the same cone –, while the majority of green ROIs clustered at larger distances (> 10 µm). The distribution of green ROIs appeared to be periodic with the average distance approximating that between cone axon terminals (approx. 8 µm, *cf.* Fig. 1C,D), indicating that these (green) ROIs were likely driven by other cones.

### HC dendritic processes “inherit” properties of the presynaptic cone

If HC dendritic tips reflect the local cone output, the measured signals are expected to share properties with the cone signals (see also Suppl. Material, Suppl. Information Fig. 1). To test this, we presented a coloured noise stimulus (Methods) and measured correlations between neighbouring cone axon terminals, and between neighbouring HC dendritic tips in the dorsal retina (Fig. 4). If HCs integrated signals globally – e.g. by averaging across a HC's dendritic arbour or by electrical HC coupling –, we would expect a higher correlation between HC dendritic tips for the two stimulus classes, due to the lateral signal spread, than for cone axon terminals. The cone recordings were performed in HR2.1:TN-XL mice [21] in which cones express the Ca^2+^ biosensor TN-XL (cf. Fig. 1D).

**Figure 4.**
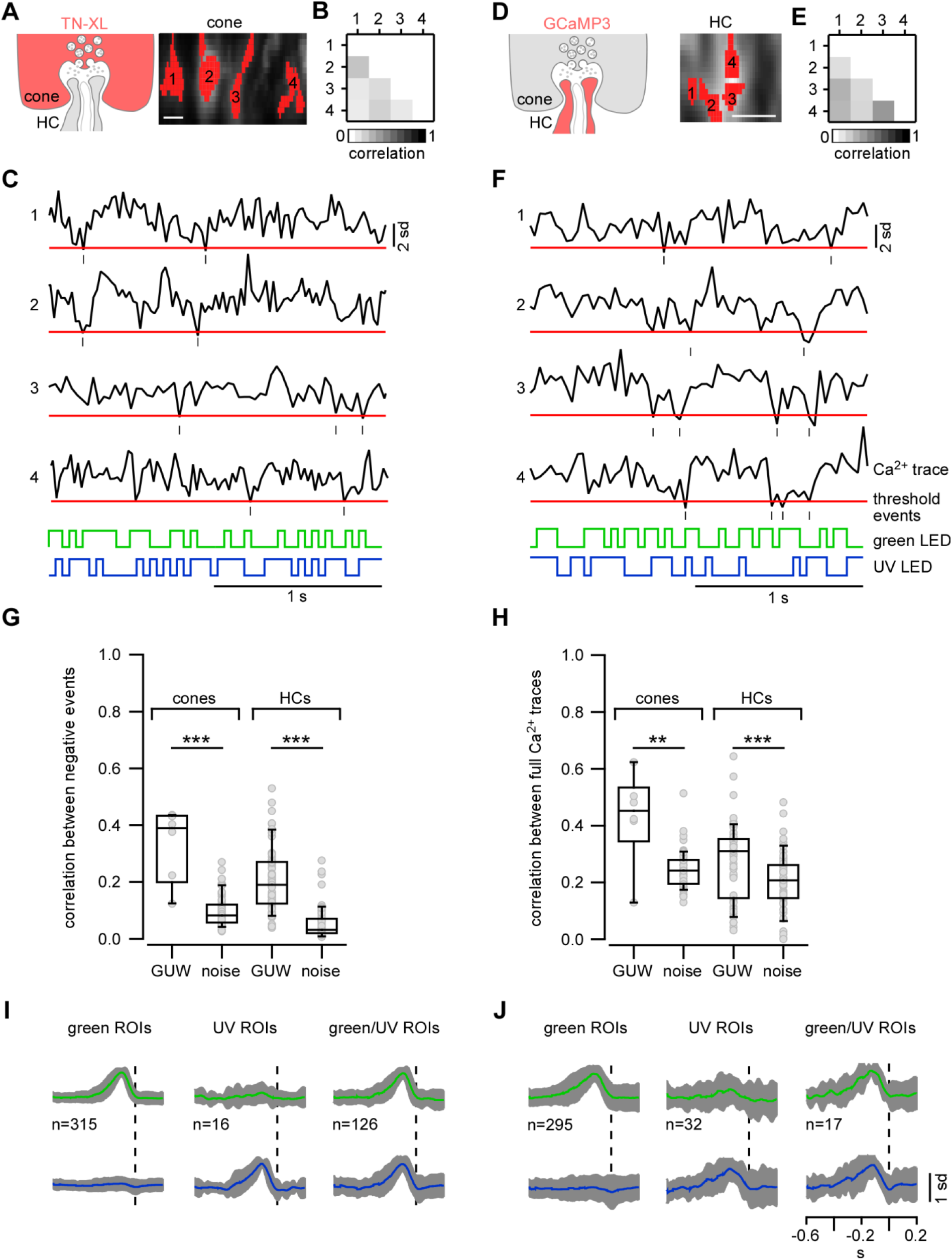
In the dorsal retina, light-evoked Ca^2+^ signals in neighbouring cone axon terminals and neighbouring HC dendrites show similar degrees of decorrelation. **A-F.** Exemplary neighbouring cone axon terminals in the HR2.1:TN-XL retina (A) and HC dendritic processes in the Cx57+^/cre^ x Ai38 retina (D) with respective Ca^2+^ signals (C, F) in response to 25-Hz coloured noise (Methods) with threshold (red line) used to detect negative events. Correlation of events for exemplary cones and HCs are shown in B and E. **G,H.** Average correlation per field for events only (G) and full Ca^2+^ traces (H) for cones and HCs in response to green, UV and white (GUW) (cones: n=6 fields; HCs: n=60 fields) and to coloured noise (cones: n=65 fields; HCs: n=57 fields). I,J. Normalized time kernels of green ROIs (amplitude green kernel > 2 s.d. noise, left), UV ROIs (amplitude UV kernel > 2 s.d. noise; middle) and mixed ROIs (amplitude green and UV kernel > 2 s.d. noise; right) for cones (I) and HCs (J) (with 2 s.d. in grey). **, p < 0.01; ***, p < 0.001. Scale bars, 5 µm.

We calculated the linear correlation coefficient (*ρ*) between Ca^2+^ traces from cone ROIs (Fig. 4A-C) in the same recording field, in response to coloured noise and to the GUW stimulus. Because the noise is a weaker stimulus compared to the GUW flashes, the correlation between cone terminal responses significantly decreased for the noise (Table 3), both when only considering negative transients (Fig. 4G) and when comparing whole traces (Fig. 4H). We repeated this experiment on HCs in Cx57^+/cre^ x Ai38 mice (Fig. 4D-F) and indeed, like for the cones, the correlation between HC responses decreased for coloured noise compared to GUW stimulation (Table 3), for negative transients (Fig. 4G) and whole traces (Fig. 4H).

**Table 3.** Linear correlation coefficient between Ca2+ traces from cones and HCs

|  | cone axon terminals (ROIs) |  | HC distal dendrites (ROIs) |  |
| --- | --- | --- | --- | --- |
| | negative events | full $\text{Ca}^{2+}$ traces | negative events | full $\text{Ca}^{2+}$ traces |
| GUW ($\rho$)<br>median<br>[MAD] | $0.390 \pm 0.043$ | $0.453 \pm 0.044$ | $0.190 \pm 0.078$ | $0.310 \pm 0.084$ |
| noise ($\rho$)<br>median<br>[MAD] | $0.082 \pm 0.030$<br>($p=3.888 \cdot 10^{-5}$ ***) | $0.242 \pm 0.040$<br>($p=0.007$ **) | $0.032 \pm 0.020$<br>($p=1.27 \cdot 10^{-17}$ ***) | $0.207 \pm 0.059$<br>($p=0.0009$ ***) |
| Δmedian | 0.308 | 0.211 | 0.158 | 0.103 |

GUW, green, UV or “white” light flashes; $\rho$ , linear correlation coefficient; MAD, median absolute deviation. Wilcoxon signed-rank test.

A direct comparison between the two sets of experiments is complicated by several factors that influence the estimation of response correlation, including different scan rates for GUW vs. noise stimuli, different biosensors in cones vs. HCs, and different ROI sizes. Nevertheless, our finding that noise stimulation results in a similar (relative) decrease in correlation for both the pre-(cone) and the postsynaptic (HC) signal (see Δmedian in Table 3) argues in favour of relatively independent signals and possibly local processing in HC distal dendrites. This is further supported by the finding that nearby HC dendrites possibly receiving input from the same cone show higher degree of correlation (correlation between negative events vs. distance for noise: Spearman R=-0.271, p=2.28·10^-20^, n=1,125; Spearman rank correlation test; Suppl. Fig. 4).

We also used the Ca^2+^ responses to the noise stimulus to estimate the temporal receptive field (time kernels, see Methods; [33]). In cone axon terminals (Fig. 4I) and HC dendritic tips (Fig. 4J), the time kernels computed using negative Ca^2+^ transients (cf. Fig. 4C,F) displayed robust positive deflections. Grouping cone ROIs by their spectral preference (derived from their time kernels, Methods) into green, UV, and mixed revealed a fraction of ~4% UV ROIs (Fig. 4I), closely matching the fraction of S-cones in the dorsal mouse retina [23]. The averaged time kernels of the different groups looked similar for cones and HCs (Fig. 4I,J); cone kernels appeared to be slightly faster, likely due to differences in biosensor properties (TN-XL: τ_decay_=0.2 s, K_D_ *in vitro=2.2* µM, from [34]; GCaMP3: τ_decay_=0.23 s, K_D_ *in vitro=0.66* µM, from [20,35]). HC kernels were noisier than those of cones. This may be related to differences in ROI area (cones, 9.6 ± 0.2 µm^2^, n=457 ROIs; HCs, 1.9 ± 0.1 µm^2^, n=344 ROIs) and, thus, different spatial averaging. The fact that we observed UV-selective kernels in HC dendritic tips just as in cones adds further evidence to the notion that HC dendritic tips show highly local Ca^2+^ signals (*cf.* Fig. 3).

### HC morphology supports electrical isolation of distal HC dendrites

To test if the HC morphology supports electrically isolating its dendritic tips, we built a simple, biophysically realistic model of a dendritic branch including synapses with cones based on a volume-reconstruction from EM data (Fig. 5; see Methods for details). First, we stimulated a single HC dendritic tip by injecting a current at the position of its synaptic cone contact such that the tip depolarized to -25 mV. We measured the resulting voltage and Ca^2+^ levels in all other cone-contacting tips and found the membrane voltage dropping rapidly with distance from the stimulated tip (Fig. 5B). Only in directly neighbouring tips (≤ 15 µm distance), the depolarization was sufficient to activate VGCCs, however, even then the resulting Ca^2+^ increase was two orders of magnitude smaller compared to the stimulated tip (Fig. 5C). We also traced the voltage change from a stimulated tip to the soma (Fig. 5D,E), showing that the tips are well isolated from the rest of the HC's dendritic arbour. Finally, we tested if our model could reproduce the light-evoked Ca^2+^ signals: We connected a representative mixture of S- and M-cones to the HC and presented full-field “light” flashes (Fig. 5F). The simulated voltage responses resembled the Ca^2+^ signals we observed in distal HC dendrites in terms of time course and spectral preference diversity (e.g. Fig. 3F). Therefore, these modelling data are in line with our experimental data, indicating that the HC dendritic morphology supports electrical isolation of distal tips and, thereby, local signalling.

**Figure 5.**
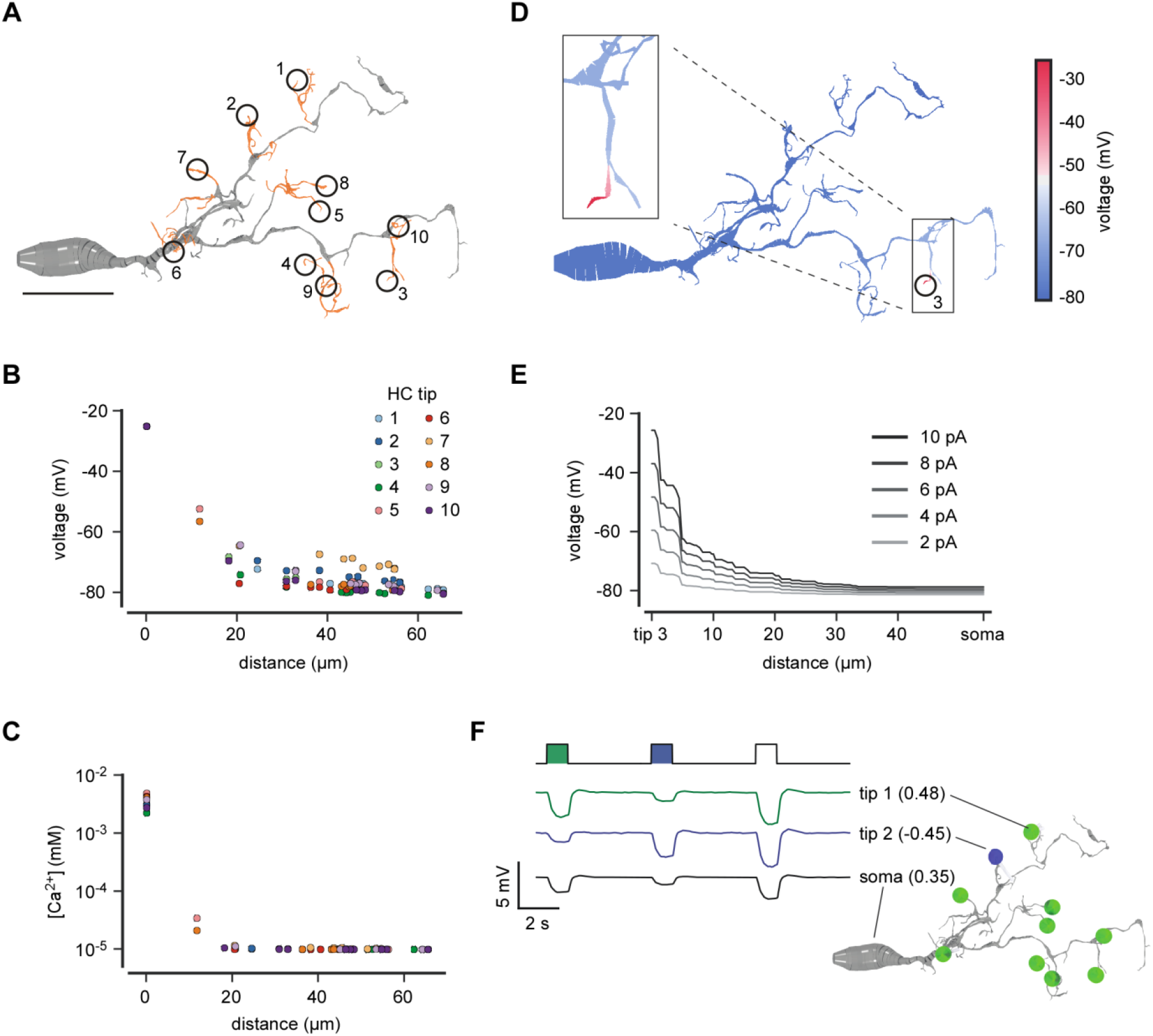
Dendritic morphology of HCs supports electrical isolation of dendritic tips. **A.** Reconstruction of a HC dendrite including soma; distal tips (orange) that invaginate cone axon terminals are numbered (1-10). Dataset from [54]. **B,C.** Voltage (B) and resulting Ca^2+^ level (C) measured in all cone-contacting HC tips as a function of distance from the current-injected tip (for details, see text). **D.** Voltage distribution across the branch after current injection into HC tip 3. **E.** Voltage drop along the dendrite between injected tip (from D) and soma. **F.** Simulated voltage responses to green/UV/white light flashes measured in two HC tips (1 and 2,contacting a green and a UV cone, respectively) and the soma. Inset: branch with connected cones (coloured by spectral preference, which is also given in brackets). Scale bar in A, 10 µm.

### Local HC feedback may shape temporal properties of cone responses

Finally, we assessed the effect of local HC feedback on the cone response. We presented a 60 Hz full-field binary noise stimulus to slices prepared from HR2.1:TN-XL mice (Fig. 6A,C) and iGluSnFR-transduced C57BL/6J mice (Fig. 6B,D) (Methods; [36]). We estimated time kernels of Ca^2+^ signals in cones and glutamate signals in the OPL as described above (cf. Fig. 4). The averaged time kernels were more transient for iGluSnFR in comparison to those for Ca^2+^ (Fig. 6E), likely reflecting differences in signal (Ca^2+^ vs. glutamate) and biosensor kinetics (τ_decay_≈ 200 ms for TN-XL vs. 92 ms for iGluSnFR, [22]). For further analysis, we computed the periodograms of the time kernels using discrete Fourier transforms [14] and examined the difference in their power spectral density for each frequency components (Methods and Fig. 6F). We first performed two consecutive recordings with an interval of 5 minutes as controls. No significant differences were found between controls for time kernels from both cone Ca^2+^ (n=61 ROIs, 11 slices, 3 mice) and glutamate release (n=76/15/3). Next, we deprived HCs from their input by bath application of NBQX and assessed the effect on the time kernels and their corresponding periodograms for cone Ca^2+^ (n=48/15/3) and glutamate release (n=47/18/3). Although the time kernels looked narrower after NBQX application, no significant differences were found between these kernels with respect to time-to-peak and *F_Area_*. However, the analysis of the periodograms revealed a significant reduction of the power spectral density at low frequencies (cone Ca^2+^, at 1 Hz, p=3·10^-4^, dependent samples t-test; glutamate release, at 0 Hz, p=3.2·10^-7^, at 1 Hz, p=4.7·10^-5^), indicating that local HC feedback contributes to temporal shaping of cone output by increasing the sensitivity of the cone signal to low frequency signal components.

**Figure 6.**
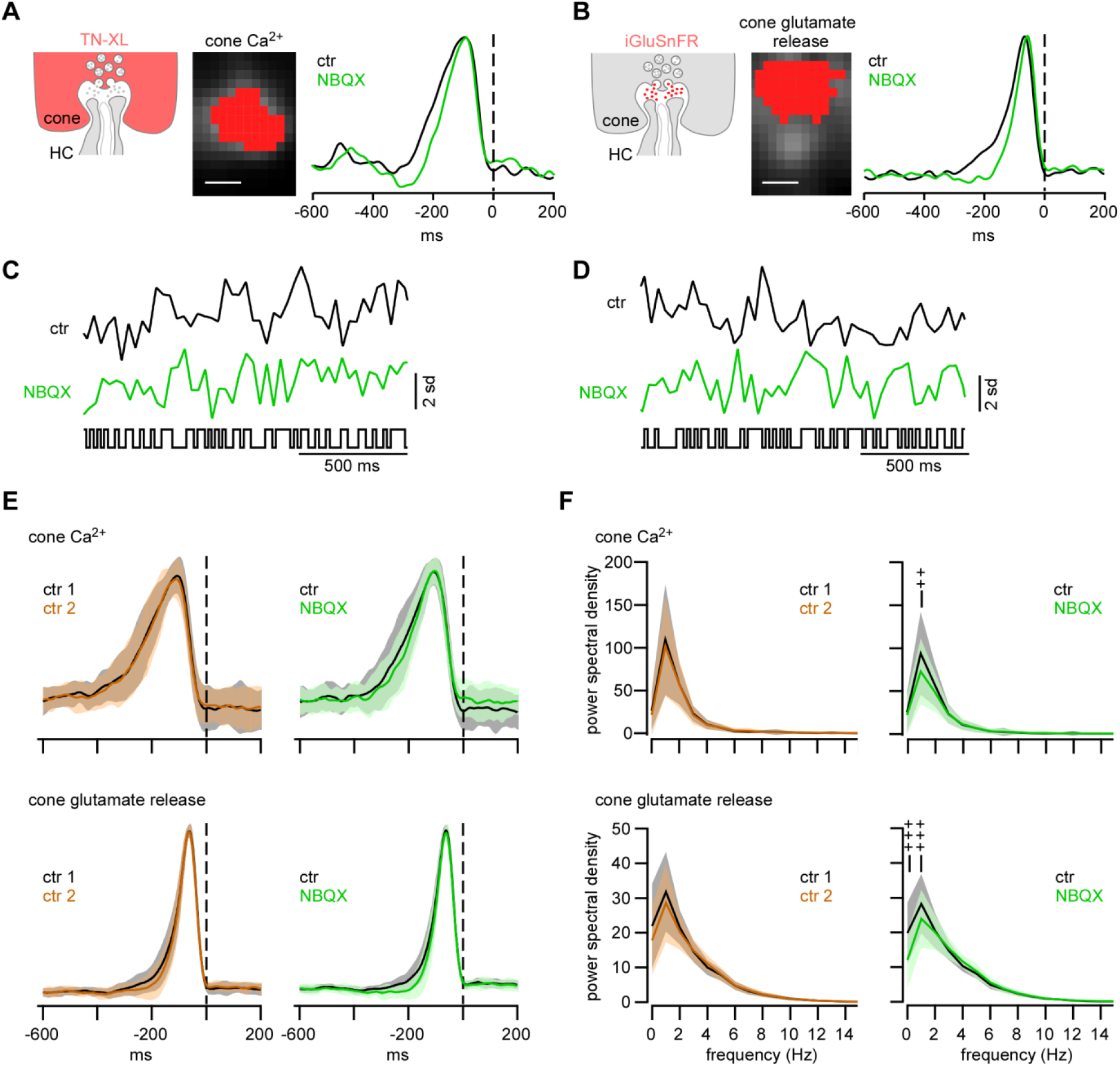
Local HC feedback modulates temporal properties of cone response. **A-D.** Exemplary ROIs of cone axon terminals, defined by TN-XL expression (A) or by iGluSnFR activity (B, Methods), with respective temporal receptive field kernels calculated from response to a full-field 60-Hz binary noise stimulus (raw traces in C, D; Methods) for control condition (black traces in A-D) and during bath application of NBQX (green traces in A-D). **E.** Normalized time kernels for cone Ca^2+^ (upper panel) and glutamate release (lower panel) for control condition (ctr1, ctr2; left) and with NBQX (right) (cone Ca^2+^: ctr, n=61 ROIs; NBQX, n=48 ROIs; cone glutamate release: ctr, n=76 ROIs; NBQX, n=47 ROIs; shaded areas indicate 1 s.d.). **F.** Periodograms (Methods) generated from cone kernels (E) using a discrete Fourier transform: cone Ca^2+^ (upper panel) and glutamate release (lower panel) for control condition (left) and with NBQX (right) (shaded areas indicate 1 s.d.). ++, p ≤ 6.66·10^-4^; +++, p ≤ 6.66·10^-5^ (Bonferroni corrected significance threshold). Scale bars, 2.5 µm.

## Discussion

Because of their large dendritic fields and gap-junctional coupling, HCs are “traditionally” thought to play a role in global processing and to provide lateral inhibition, e.g. for contrast enhancement, in the outer retina (reviewed in [19]). Recent studies, however, suggest the existence of a local processing mode, in which HCs provide locally “tailored” feedback to individual cones [14,18] - reminiscent of the local dendritic processing in amacrine cells (e.g. [4]).

Here, we recorded light stimulus-evoked pre- and postsynaptic activity at the cone-HC synapse in the mouse retina and present three lines of evidence supporting that mouse HCs can process cone input in a highly local and independent manner: First, neighbouring dendritic tips, which presumably contact neighbouring cones, differed in their chromatic preferences. While the ubiquitous GCaMP3 expression in HCs did not allow us to assign ROIs to individual HCs, it is unlikely that our data are solely explained by recording two overlapping “kinds” of HCs with opposite spectral preference, simply because mice feature only one type of HC, which indiscriminately contacts all cones within its dendritic field [29,37]. Second, the correlation levels of Ca^2+^ signals measured in neighbouring HC dendritic tips were similar to those recorded in neighbouring cone axon terminals. If cone inputs were already averaged at the level of the distal HC dendrite, we would have expected an increase in correlation from cones to HCs. Hence, our correlation data supports local signalling (and possibly processing) in HC dendritic tips. Third, a simple, biophysically realistic model confirms that the HC morphology supports electrical isolation between dendritic tips.

By isolating HCs pharmacologically from their cone input, we showed that the HC feedback may shape the temporal filtering properties of the cone synapse, i.e. by modulating the power at low stimulus frequencies. Taken together, our study extends the “traditional” view of global HC signalling by a powerful local component, indicating that dendritic processing already happens at the first synapse of the retina.

### Local vs. global HC feedback

Byzov and Shura-Bura [38] were the first to suggest that HCs provide local feedback to cones. Following experimental confirmation, it was then proposed that both local and global feedback are triggered by the activation of AMPA/KA receptors on HCs [18], but with local feedback being mediated by the local Ca^2+^ in the dendritic tip, and global feedback relying on depolarisation and possibly amplification by VGCCs [39]. While we did not find interactions between distal HC dendrites, the “mixing” of S-and M-signals we observed in proximal HC dendrites hints at some degree of global signal integration. We cannot exclude the possibility that the slice preparation introduces a bias towards local signalling, however, earlier work in rabbit HCs suggests that local feedback suffers more from slicing than global feedback [18]. Furthermore, more global-scale interactions have been successfully demonstrated in mouse retinal slices; these include, for instance, lateral inhibition between cones [12] and electrical coupling within the AII amacrine cell network [40]. Global signal integration within the HC network requires Cx57-mediated gap-junction coupling [17]. In the Cx57^+/cre^ x Ai38 mice used here, one *Cx57* allele is replaced by a *cre* gene, resulting in a reduced Cx57 expression. The HCs in these mice feature smaller receptive fields (RFs) and elevated resting potentials, but since HC coupling is intact and cone-HC synapses seem unaltered [41], we do not expect this genetic modification to substantially affect our conclusions.

### Mechanism(s) of local Ca^2+^ signalling in HC dendrites

What is the cellular basis of the local Ca^2+^ signalling we observed in HC dendrites? In line with previous studies [28,29,31], we show that these signals are mediated by a combination of Ca^2+^-permeable AMPA/KA-type glutamate receptors, VGCCs, and Ca^2+^ released from stores. This combination is reminiscent of another reciprocal synapse with local signalling: The synapse between rod bipolar cells (RBCs) and A17 amacrine cells [42]. Here, Ca^2+^ enters a dendritic AC varicosity via AMPA receptors and triggers GABA release, with the necessary amplification of the Ca^2+^ signal generated by Ca^2+^-induced Ca^2+^ release from stores. To keep the signal from spreading to neighbouring varicosities, A17 cells express Ca^2+^-activated potassium (BK) channels that hyperpolarize the varicosity and suppress activation of VGCCs. In addition, varicosities are spaced with an average distance of ~20 µm along the dendrite to increase electrical isolation.

Local signalling in HCs may employ a similar mechanism: (*i*) As shown for several species, local HC feedback can be triggered by AMPA receptor activation without requiring VGCCs [18]. If this is also true for mouse HCs, is still unclear. The Ca^2+^ signals evoked by AMPA/KA puffs mainly involved L-type VGCCs and Ca^2+^ stores, but for weaker, more physiological stimuli (i.e. light), the direct contribution of AMPA/KA receptors to the Ca^2+^ signals may be greater, as it seems to be the case in A17 cells [42]. Moreover, activity of VGCCs in HCs is suppressed by dopamine, which is released in a light-dependent fashion [43], suggesting that VGCCs contribute less to the Ca^2+^ signal under our light conditions. (*ii*) HCs express BK channels that limit membrane depolarisation in a voltage - and Ca^2+^-dependent manner [44]. (*iii*) Ca^2+^ signals in HC dendrites partially rely on Ca^2+^ stores [28]. (*iv*) The HC morphology enhances electrical isolation between dendritic tips, as supported by our modelling data.

### Do rods contribute to the Ca^2+^ signals in HC dendrites?

The transgenic mouse expresses GCaMP3 in all HC compartments, and because dendritic and axonal HC processes are intermingled, we could not distinguish them solely based on their morphological appearance. Yet, our conclusions rely on the assumption that the measured Ca^2+^ signals reflect cone input and that rod input (either mediated by direct rod-HC contacts or by rod-cone coupling) can be neglected. We think that this was the case for three reasons: (*i*) We measured the largest Ca^2+^ signals at the OPL level where HC dendrites invaginate the cone axon terminal base [19,27]. (*ii*) The chromatic tuning of these Ca^2+^ signals reflected the local ratio of S-vs. M-opsin expression along the retina's dorso-ventral axis. If rods had substantially responded to either UV or green, we would have expected an additional UV response in dorsal HCs and/or an additional green response in ventral HCs. (*iii*) Laser-evoked photoreceptor excitation alone generated a “background illumination” equivalent to ~10^4^ P*·s^-1^/cone (Methods; [32]), which is probably similar in rods [25]. Electrical recordings from mouse rods in slices indicate that rod photoresponses disappear at ~10^4^ P*·s^-1^/rod [45], suggesting that rods were not operational under our stimulation conditions. Our earlier finding that RBCs respond to light-on stimuli under similar conditions [36] may be explained by direct (or indirect) cone input to RBCs, as in mice, ~70% of the RBCs contact at least one cone [46], and cones and rods are at least weakly coupled at our light levels [47].

### Functional consequences of local dendritic processing for HC feedback to cones

What could be the purpose of local HC feedback to cones? Our pharmacological data indicate that (local) HC feedback boosts low frequency signals in the cone output, consistent with a general role of HC feedback in dynamically shaping/adapting the time course of cone transmission, e.g. as a function of stimulus configuration (e.g. centre vs. surround stimulation; [48,49]). Specifically, our finding is consistent with earlier work showing that activation of HCs (by a dark annular stimulus) shifts the frequency sensitivity of a HC in the stimulus center towards lower frequencies [50]. That we could not also find a shift at higher frequencies may well be due to temporal limitations both of the imaging system and the biosensors (cf. Results).

In theory, the objective of sensory neurons is often considered to be the relative enhancement of relevant information content from a sensory input, given a limited metabolic capacity [51]. Adaptational mechanisms allow the circuitry to robustly meet this objective despite changing natural scene statistics [52], whether by enhancing features (increasing information content) or removing redundancy (reducing metabolic cost). For HCs, these elements have typically been considered for adaptation to spatial properties, through mechanisms such as the centre-surround RF and background-luminance subtraction. Here, the local adaptation observed appears to operate in the time domain; the effect is made visible by changes in sensitivity to (at least but probably not solely) low frequency components, which might convey little information content in natural scenes where they are strongly present, and more information otherwise. In this sense, local feedback may serve temporal contrast enhancement. That the feedback can occur at the level of individual photoreceptors is perhaps not surprising, as such a cell driven in isolation is still subject to trade-offs in information and metabolic cost. In natural scenes, spatial and temporal statistics are not independent, and the interplay between their respective adaptational mechanisms – such as here the potential interplay between local and global feedback mechanisms in HCs – is an attractive subject for further study (see discussion in [19]). A promising approach to tackle these questions may be a combination of voltage biosensors [53] to probe the voltage “distribution” across an HC's dendritic arbour with biophysically realistic models.

## Author Contributions

C.A.C., T.S. and T.E. designed the study; C.A.C. and S.P. established the experimental approach; C.A.C. performed experiments and pre-processing; C.A.C. analysed the data, with input from T.S., T.B., L.E.R., P.B., and T.E.; C.B. generated and analysed the model, with input from T.S., P.B., and T.E.; C.A.C., T.S., T.E., and L.E.R., wrote the manuscript, with input from P.B., C.B. and T.B.

## Acknowledgements

This work was supported by the Deutsche Forschungsgemeinschaft (EXC 307, CIN to T.E. and T.S.; SCHU 2243/3-1 to T.S., BE5601/1-1 to PB) and the German Ministry of Science and Education

(BMBF) through the Bernstein Award to PB (FKZ: 01GQ1601). We thank G. Eske for excellent technical assistance and K. Franke for performing the intravitreal virus injection; Y. Zhang for tracing the HC dendrite and R. G. Smith for help in setting up the NeuronC model. We thank L.L. Looger, the Janelia Research Campus of the Howard Hughes Medical Institute and the Genetically-Encoded Neuronal Indicator and Effector (GENIE) Project for making the viral construct AAV9.hSyn.iGluSnFR.WPRE.SV40 publicly available.

The authors declare no conflict of interest.

---

**Supplementary Figure 1.**
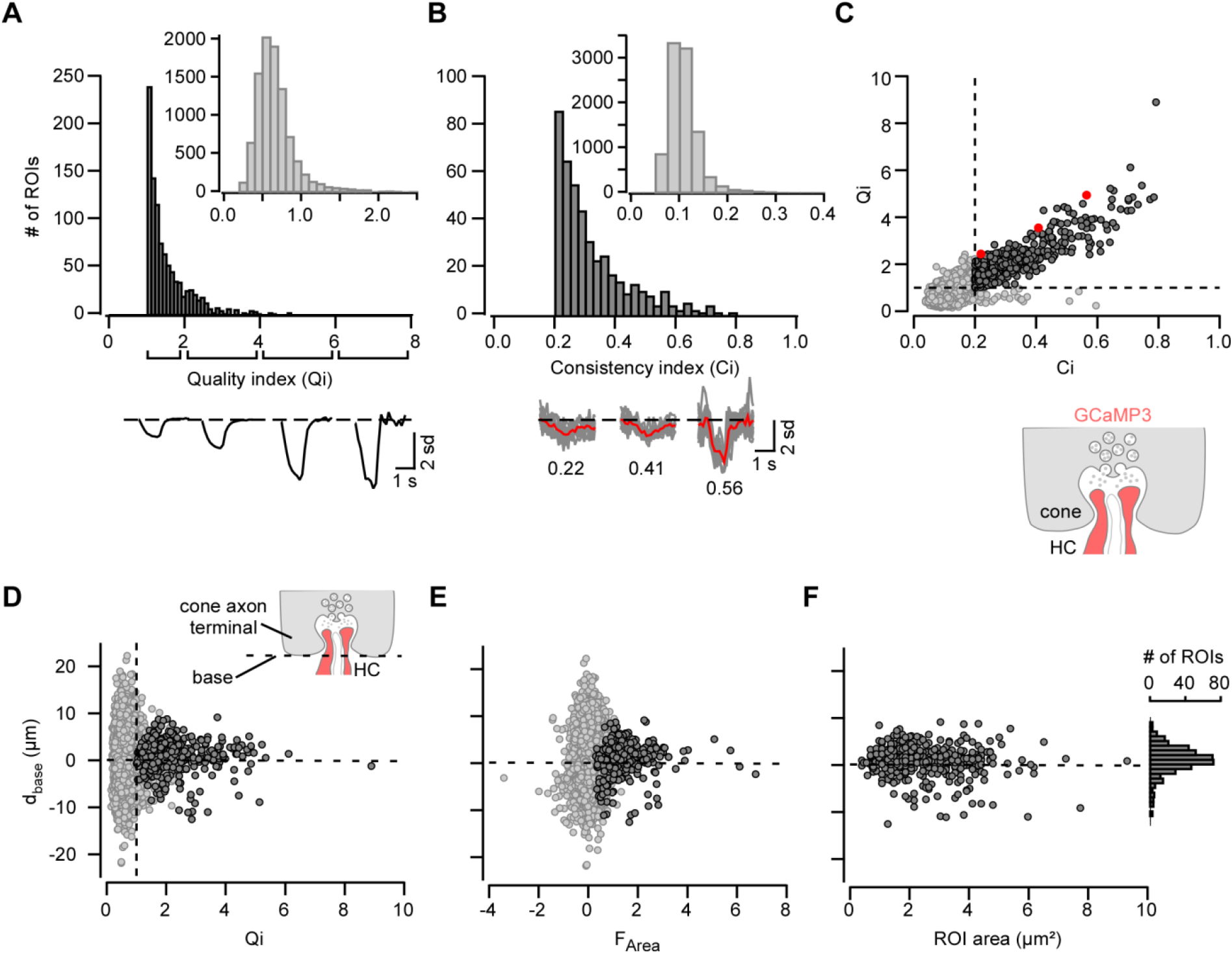
Selection of ROIs on horizontal cell processes based on their light-evoked Ca^2+^ signals. **A.** Distribution of quality index (*Qi*), defined as ratio between Ca^2+^ response amplitude to white flash and s.d. of noise (Methods). Only ROIs with *Qi* ≥ 1 (dark grey) were considered for further analysis (inset shows distribution of discarded ROIs). *Below:* Average Ca^2+^ responses across ROIs for different *Qi* intervals. **B.** Distribution of consistency index (Ci), defined as ratio between variance of the mean and mean of variance (Methods). Only ROIs with *Ci* ≥ 0.2 (dark grey) were considered for further analysis (inset shows distribution of discarded ROIs). *Below:* Exemplary Ca^2+^ traces for different *Ci* values (mean in red, n=10 trials in grey). **C.** *Qi* as a function of *Ci*, with ROIs passing both criteria shown as dark-grey dots (n=423 of 9,912 ROIs passed both criteria). Red dots indicate ROIs of example traces in B. **D,F.** Distance of ROI (centre of mass) to cone axon terminal base (*d_base_*) as a function of *Qi* (D, the dashed line indicates the cone base), area-under-the-curve (*F_Area_*, E) and ROI area (F). ROI areas (F) above the cone axon terminal base (*d_base_*> 0; 2.45 ± 0.07 µm^2^, n=292) and below (*d_base_* < 0; 2.87 ± 0.30 µm^2^, n=131) were not significantly different (p=0.248). Histogram shows distribution of ROIs along the cone terminal base. Diagram on the right depicts imaged synaptic compartment and used biosensor (red).

**Supplementary Figure 2.**
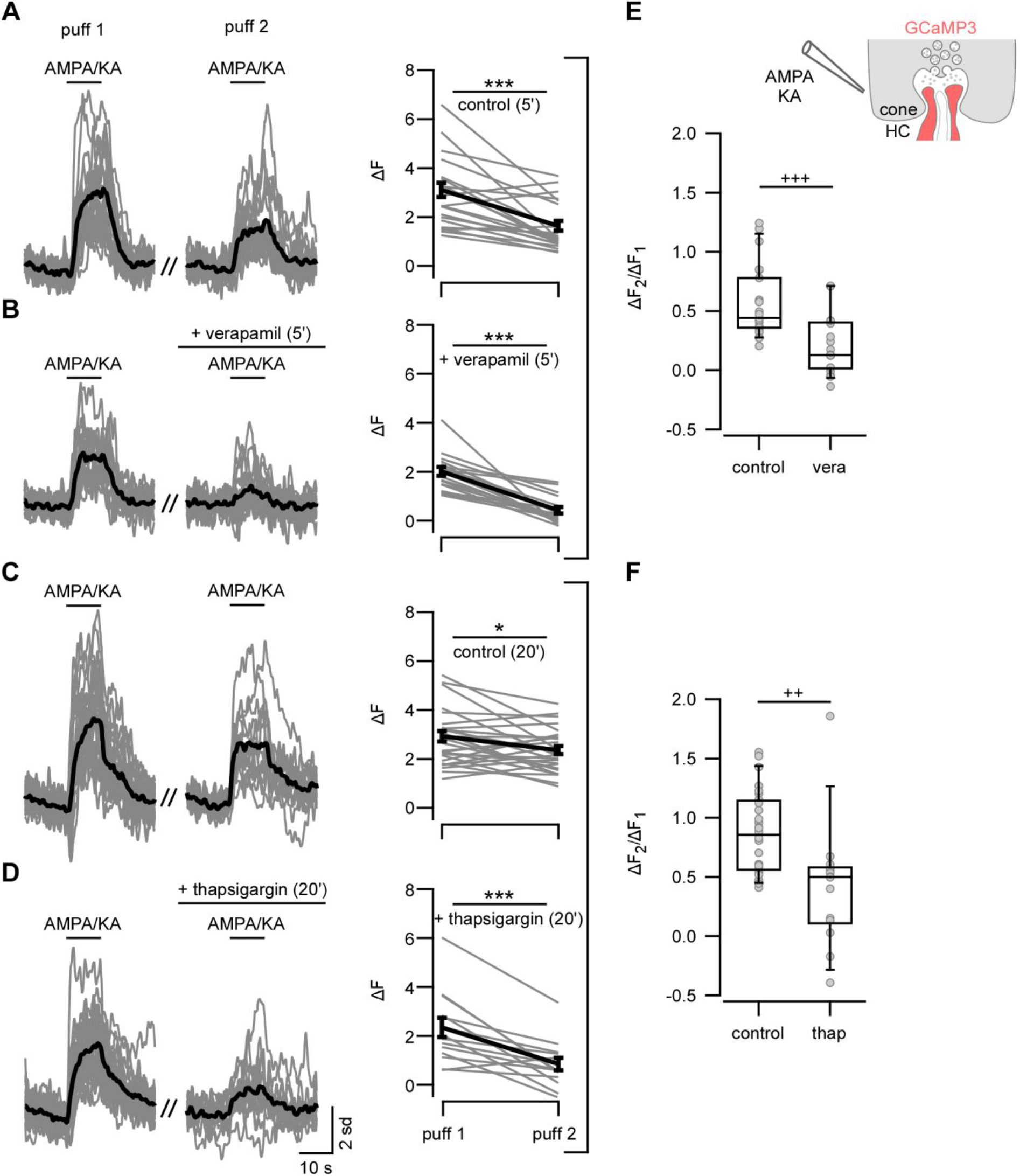
Ca^2+^ signals in HC dendrites are mediated by voltage-gated Ca^2+^ channels and intracellular Ca^2+^ stores. **A-D.** Ca^2+^ signals in HC processes evoked by two consecutive AMPA/KA puffs (short bars indicate puff timing). *Left column:* in standard bathing medium; *middle column:* normal medium for 5 min (A, n=23 ROIs from 2 slices, 2 mice), with bath application of verapamil for 5 min (B, n=18, 3 slices, 2 mice), normal medium for 20 min (C, n=28, 3 slices, 2 mice), and with bath application of thapsigargin for 20 min (D, n=13, 5 slices, 3 mice). *Right column:* Quantification of drug effects on response amplitude *ΔF* (error bars indicate SEM; *, p < 0.05; ***, p < 0.001). **E,F.** Ratios between *ΔF*_2_ (2^nd^ puff) and *ΔF*1 (1^st^ puff) control, verapamil (vera) (E) and thapsigargin (thap) (F) (++, p ≤ 0.005; +++, p ≤ 0.0005) (Bonferroni corrected significance threshold).

**Supplementary Figure 3.**
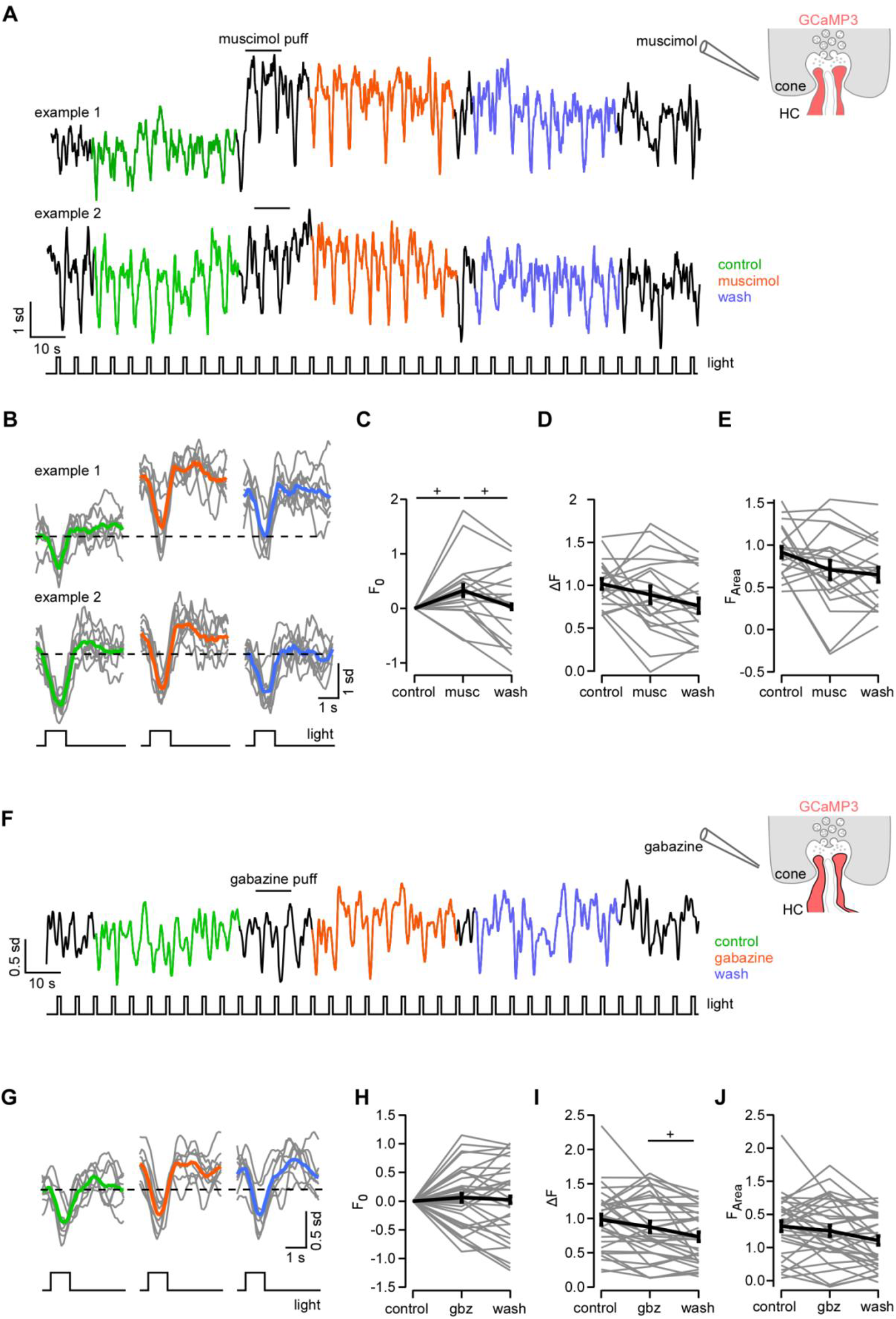
GABA modulates light-evoked Ca^2+^ signals in HC dendrites. **A.** Two exemplary Ca^2+^ responses of HC processes to white flashes before (control), after a puff of the GABAa receptor agonist muscimol and during wash-out illustrating the variability of the response to the muscimol puff. **B.** Averaged responses for control (green), muscimol (orange) and wash-out (blue) for the two exemplary traces shown in (A). **C-E.** Quantification of muscimol (musc) effects on response baseline (*F_O_*, C), amplitude (ΔF, D) and area-under-the-curve (*F_Area_*, E) (average of n=20 ROIs from 4 slices, 2 mice). **F.** Experiment as in (A) but for the GABA_A_ receptor antagonist gabazine. **G.** Averaged responses for control, gabazine and wash-out. **H-J.** Quantification of gabazine (gbz) effects (average of n=33 ROIs, 4 slices, 2 mice). Error bars indicate SEM. +, p ≤ 0.025 (Bonferroni corrected significance threshold).

**Supplementary Figure 4.**
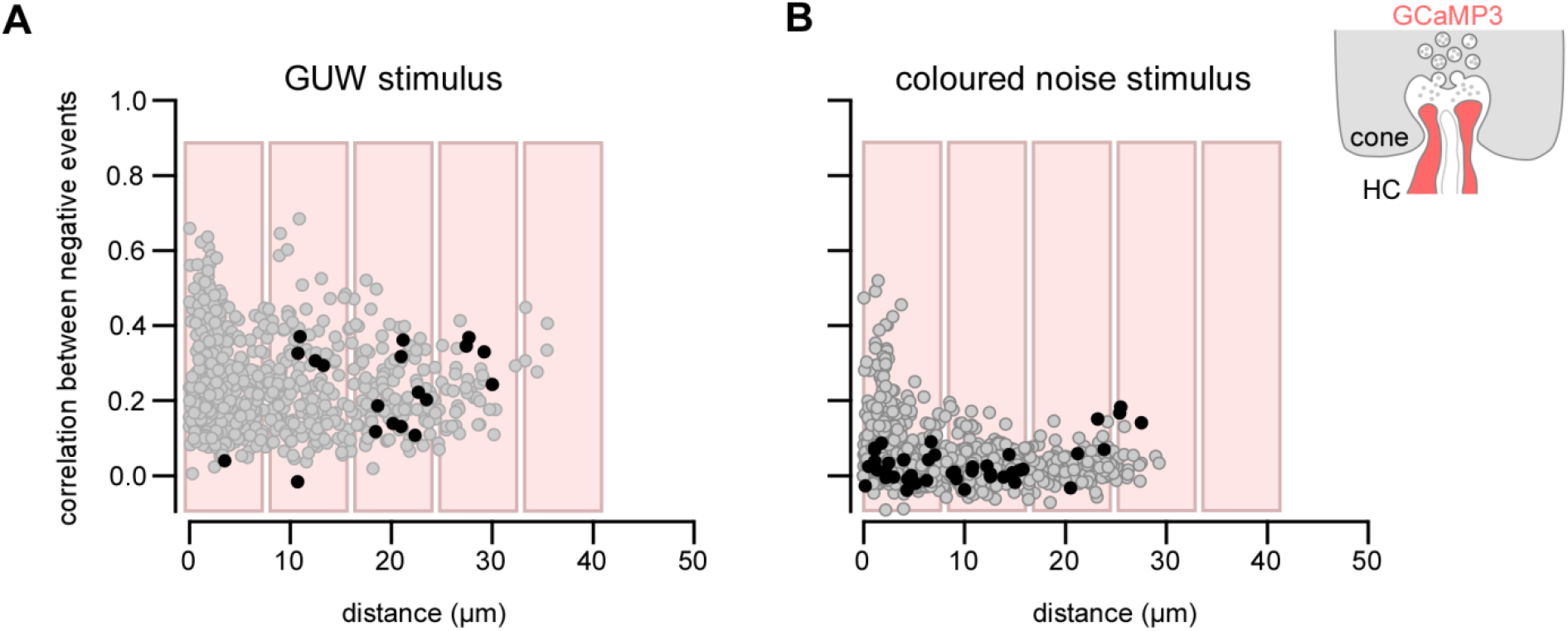
Correlation between negative events in HC processes. **A,B.** Correlation between negative events in HC processes (Cx57^+/cre^ x Ai38 mouse line) for all combinations of ROIs present in the same field as a function of their distance; for GUW stimulus (A) and coloured noise (B). Dark circles indicate the combinations between “purely” green and “purely UV” ROIs (GUW, *|SC|* > 0.4; coloured noise, amplitude UV or green kernel > 2 s.d. noise). Reddish boxes illustrate expected location of neighbouring cone axon terminals.

## STAR Methods

### Animals

For the Ca^2+^ imaging experiments in retinal horizontal cells (HCs), we crossed the transgenic mouse lines Cx57^cre/cre^ [31] and *B6;129S-Gt(ROSA)26Sor^tm38(CAG-GCaMP3)Hze^/J* (Ai38) [20], yielding Cx57^+/cre^ x Ai38 mice, which express the Ca^2+^ biosensor GCaMP3 [35] selectively in HCs. For Ca^2+^ imaging in cone axon terminals, we used the HR2.1:TN-XL mouse line [21], which expresses the FRET-based Ca^2+^ biosensor TN-XL [55] exclusively in cones. TN-XL enables ratiometric Ca^2+^ measurements by determining the ratio between the signals of its two fluorophores, eCFP (FRET donor) and citrine (FRET acceptor), which are linked by the Ca^2+^ sensor troponin C (for details, see [21]). For glutamate imaging, iGluSnFR [22] was ubiquitously expressed in the retina after intra-vitreal virus injection in C57BL/6J mice (see Virus injection). We observed iGluSnFR expression predominantly in HCs, likely because bipolar cells express AAV constructs less efficiently [56]. Both male and female adult mice (4-18 weeks of age) were used. Animals were deeply anesthetized with isoflurane (CP-Pharma, Germany) and killed by cervical dislocation. All procedures were performed in accordance with the law on animal protection (Tierschutzgesetz) issued by the German Federal Government and approved by the institutional committee on animal experimentation of the University of Tübingen.

### Retinal tissue preparation

For all imaging experiments, mice were dark adapted for at least 2 hours and then killed. Under dim red light, both eyes were marked at the ventral side to maintain retinal orientation, quickly enucleated and hemisected in carboxygenated (95% O2 / 5% CO2) extracellular solution with (in mM): 125 NaCl, 2.5 KCl, 1 MgCl2, 1.25 NaHCO3, 20 glucose, 2 CaCl2, 0.5 L-glutamine and 150 µM pyridoxal 5-phosphate (a cofactor of the glutamic acid decarboxylase, [57]) (Sigma-Aldrich or Merck, Germany). Cornea, lens and vitreous body were carefully removed. The retina was separated from the eye-cup, cut in half, flattened and mounted photoreceptor side-up on a nitrocellulose membrane (0.8 µm pore size, Millipore, Ireland). Using a custom-made slicer [58], acute vertical slices (300 µm thick) were cut parallel to the naso-temporal axis. Slices attached to filter paper were transferred on individual glass coverslips, fixed using high vacuum grease and kept in a storing chamber at room temperature for later use. For all imaging experiments, individual retinal slices were transferred to the recording chamber, where they were continuously perfused with warmed (~36°C), carboxygenated extracellular solution containing 0.5 µM sulforhodamine 101 (SR101; Sigma-Aldrich, Germany) to visualize cone axon terminals.

### Virus injection

Before the injection of AAV9.hSyn.iGluSnFR.WPRE.SV40 (Penn Vector Core, PA, USA), mice (5-7 weeks) were anaesthetized with 10% ketamine (Bela-Pharm GmbH, Germany) and 2% xylazine (Rompun, Bayer Vital GmbH, Germany) in 0.9% NaCl (Fresenius, Germany). A Hamilton syringe (syringe: 7634-01, needle: 207434, point style 3, length 51 mm, Hamilton Messtechnik GmbH) containing the virus was fixed on a micromanipulator (M3301, World Precision Instruments, Germany) at an angle of 15°. Then, 1 µl of the virus was injected into the naso-ventral part of the vitreous body [36]. Recordings were performed 3 weeks after the injection.

### Two-photon imaging

Ca^2+^ and glutamate signals were recorded on a customized MOM-type two-photon microscope (Sutter Instruments, Novato, CA; designed by W. Denk, MPI for Neurobiology, Martinsried, Germany) [25,59], equipped with a mode-locked Ti:Sapphire laser (MaiTai-HP DeepSee; Newport Spectra-Physics, Germany) tuned to either 860 or 927 nm for TN-XL and GCaMP3/iGluSnFR excitation, respectively, and a 20x water-immersion objective (XLUMPlanFL, 0.95 NA, Olympus, Germany, or W Plan-Apochromat 20x/1.0 DIC M27, Zeiss, Germany). Two PMTs with appropriate band-pass filters were used to detect the fluorescence emission of (a) TN-XL/citrine, GCaMP3 (538 BP 50, AHF, Germany) or iGluSnFR (510 BP 84), and (b) TN-XL/eCFP (483 BP 32) or SR101 (630 BP 60). We acquired time-lapsed image series with the custom software ScanM (by M. Müller, MPI for Neurobiology, and T. Euler) running under IgorPro 6.37 (Wavemetrics, Lake Oswego, OR, USA). Images of 128 x 64 pixels (51.8 x 28.2 µm or 38.7 x 20.8 µm) at a frame rate of 7.8125 Hz were recorded for all visual stimuli except the “coloured noise” and binary noise stimuli (see below), where we used images of 128 x 16 pixels (51.8 x 7.1 µm or 38.7 x 5.2 µm, at 31.25 Hz). Recording fields were always located at the outer plexiform layer (OPL) to prevent bleaching of the cone outer segments by the scanning laser [21,32].

Because SR101 is endocytosed by terminals of synaptically active cells such as photoreceptors [26], we performed control experiments to rule out that SR101 fluorescence in the “red” channel contributed to the activity signals measured in the “green” channels. To this end, we presented light flashes to retinal slices from C57BL/6 mice bathed in SR101 and recorded the fluorescence signal in both channels. Indeed, we did not find any light stimulus-dependent signal modulation in the red channel (data not shown). Furthermore, we did not detect any substantial SR101 fluorescence in the green channel.

### Light stimulation

Full-field light stimuli were generated by two band-pass-filtered LEDs (UV, 360 BP 12; green, 578 BP 10; AHF) driven by an open-source microprocessor board (http://www.arduino.cc) and synchronized with the scanner retrace to avoid light stimulus artefacts during image acquisition. The light from the two LEDs was combined by a beam-splitter (400 CDLP, AHF) and focused on the retinal slice through the bottom of the recording chamber via a condenser lens (H DIC, 0.8 NA, Zeiss). The intensity of each LED was adjusted such that the photoisomerisation (P*) rate in S-cones elicited by the UV LED was equal to the P* rate elicited by the green LED in M-cones [60,61]. The light intensity generated by each LED was equivalent to P* rates (in P*s^-1^/cone) ranging from 0.5·10^3^ (I_min_) to 6.5·10^3^ (I_max_) for all stimuli except the binary noise stimulus (I_MIN_= 0.6·10^3^, I _MAX_=19·10^3^), where a different light stimulator was used (for details, see [36]). Note that the two-photon excitation laser caused an additional steady background illumination (Ibkg) of approx. 10^4^ P*s^-1^/cone [21,32], which likely amounts to a similar P* rate in rods [25]. Under these stimulus conditions, it is unlikely that rods are operational (see Discussion).

Note that we use the term “white” to refer to the simultaneous stimulation with both LEDs with the same P* rate. All following 5 stimulus protocols were preceded by a 15-s period that allowed the photoreceptors adapting to the background (*I_MIN_ + I_BKG_*):

a. A white flash protocol consisting of 1-s bright flashes (from a background of *I_MIN_* to *I_MAX_*) of “mouse-white” (both LEDs on) at 0.2 Hz. This protocol was used to assess drug effects on light-evoked Ca^2+^ responses.
b. A colour flash protocol consisting of bright green, UV and white 1-s flashes ("GUW") at 0.2 Hz and repeated 10 times for each colour (same intensity levels as for (a)). This protocol was used to determine the spectral contrast (SC, see below) preference.
c. A contrast and colour flash protocol consisting of 1-s bright and dark flashes, with the respective LED combinations (green, UV, and white) at *I_max_* or *I_min_*, respectively, at 0.2 Hz and repeated 8 times for each condition (Intensity between flashes: 3·10^3^ P*s^-1^/cone). This protocol was used to determine the *SC* and the dark-light index (*DLi*, see below).
d. A “coloured noise” stimulus protocol consisting of a 25 Hz pseudo-random sequence of green, UV, white, and dark flashes. This protocol was used to probe correlation between neighbouring cones and HC processes and to calculate time kernels (see below).
e. A binary noise stimulus protocol consisting of a 60-Hz pseudo-random sequence of dark and bright flashes. This protocol was also used to calculate time kernels.

### Immunohistochemistry

After two-photon imaging, a subset of retinal slices were fixed with 4% paraformaldehyde (PFA) in 0.1 M phosphate-buffered saline (PBS) at 4°C for 15 min. Slices were then washed in 0.1 M PBS, and submerged in blocking solution (0.1 M PBS, 0.3% Triton X-100, 10% donkey serum) over night at 4°C. Afterwards, slices were incubated for 4 days at 4°C with primary antibodies (rabbit anti-M-opsin (1:1,000) from EMD Millipore, Billerica, MA, USA; goat anti-S-opsin (1:500) from Santa Cruz Biotechnology (Germany) in 0.1 M PBS, 0.3 Triton X-100, and 5% donkey serum. The following day, slices were washed in 0.1 M PBS and incubated with the secondary antibodies (donkey anti-rabbit conjugated to Alexa Fluor 568 (1:1000) and donkey anti-goat conjugated to Alexa Fluor 660 (1:1000), both Invitrogen, Carlsbad, CA, USA). Image stacks (15 frames of 1024 x 1024 pixels, 15 µm Z-steps) were acquired on a confocal laser-scanning microscope (Leica TCS SP8, Germany) which was equipped with green (552 nm) and far-red (638 nm) lasers and a 10x 0.3 NA objective lens (Leica). Maximum-intensity projections of the image stacks were performed using Fiji (http://fiji.sc/Fiji).

### Pharmacology and drug application

All drugs were prepared as stock solutions in distilled water or, in the case of thapsigargin, in DMSO (0.1% in the extracellular medium), and were stored at -20°C. Before each experiment, drugs were freshly diluted from stock solution in carboxygenated extracellular solution. For puff application, a glass electrode (tip diameter: 1-2 µm) was placed approx. 100 µm above the recorded region of the slice and drug solution was puffed for 10 s using a pressure application system (0.2-1 bar, Sigmann Elektronik GmbH, Germany). The lateral spread of the puff was about 200 µm in diameter, as measured by puffing a fluorescent dye (SR101). For bath application, the tissue was perfused with the drug added to the bathing solution for 5 min or, in the case of thapsigargin, for 20 min (perfusion rate of ~1.5 ml/min). For puff application, the following concentrations were used (in µM): 200 6,7-dinitroquinoxaline-2,3-dione (NBQX), 50 a-amino-3-hydroxy-5-methyl-4-isoxazolepropionic acid (AMPA), 25 kainic acid (KA), 100 muscimol and 100 SR-95531 hydrobromide (gabazine). For bath application, we used (in µM): 100 verapamil, 5 thapsigargin and 100 NBQX. All drugs were purchased from Tocris Bioscience (Bristol, England) except for KA, which was purchased from Sigma-Aldrich.

### Data analysis

To analyse light-evoked Ca^2+^ signals in HCs and cones, as well as glutamate release in the OPL, we used custom-written scripts in IgorPro (Wavemetrics) and SARFIA [62]. For GCaMP3 and TN-XL fluorescence (Ca^2+^ in HCs and cones, respectively), regions-of-interest (ROIs) were anatomically defined using SARFIA's automatic Laplace operator feature on the averaged, filtered image series and manually corrected if required (e.g. if two nearby structures shared one ROI); ROIs with an area < 10 pixels were discarded. For iGluSnFR fluorescence (glutamate release), the correlation over time between neighbouring pixels was measured and ROIs were determined based on a correlation threshold (defined for each recording depending on the signal-to-noise ratio). ROI diameters were limited to range between 5 to 8 µm (diameter of a cone axon terminal).

To determine the spatial resolution of our system, we measured the point spread function (PSF) using fluorescent beads (0.17 µm in diameter; Invitrogen) and found the resolution (Abbe) limited to 0.6 µm in the x-y plane and 4.1 µm along the z axis (FWHM of Gaussian fit, green channel). Cone axon terminals measure approx. 5 µm in diameter, distal HC dendrites between 1 to 2 µm [37]. Therefore, both can be resolved in the x-y plane. Due to the limited z resolution, it is possible that a ROI averaged across multiple fine distal HC dendrites stacked along the z-axis. In case of the much larger cone axon terminals, averaging across two stacked ones is unlikely but cannot be excluded. In any case, however, since such averaging is not expected to increase functional diversity between individual ROIs, it should not have affected our conclusions.

To estimate each ROI's “vertical” position within OPL, the positions of cone axon terminals were visualized using SR101 fluorescence (Fig. 1C,D). Here, cone axon terminals can be identified because they are more brightly stained compared to rod axon terminals and HC processes, and because they are organized along the OPL like beads on a string. A ROI's distance to the cone axon terminal base (d_base_) was estimated relative to a manually drawn straight line tracing the base of all cone axon terminals in a recorded field, using the shape transition in brightness between the cone axon terminals and the weakly labelled HC dendrites below as a landmark.

For TN-XL, the ratio between acceptor (citrine) and donor fluorescence (eCFP) was calculated on the image series, prior to signal extraction. For all indicators, time traces were extracted for each ROI, de-trended by high-pass filtering at ~0.1 Hz (except for the analysis of drug effects on the baseline) and z-normalized 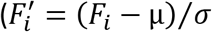, with samples F¿, and mean (µ) and s.d. (ơ) of the trace). For all flash stimuli, we determined response amplitude (ΔF), area-under-the-curve (*F_Area_*) and, in case of NBQX, muscimol and gabazine puffs, as well as for the contrast and colour flash protocol, also the Ca^2+^ baseline level (F_0_). These parameters were measured on the trace smoothed using IgorPro's boxcar algorithm with 2 passes for all stimuli (except for drug experiments, where 5 passes were used).

Two quality criteria were defined to identify responsive ROIs: The **quality index** (*Qi*) is defined as the ratio between *ΔF* in response to a white flash and the s.d. of the noise of the trace (= raw trace minus the trace smoothed using IgorPro's boxcar algorithm with 2 passes). For stimulus protocol (c), *Qi* was calculated independently for dark and bright flashes. Depending on stimulus and experiment type, we used different *Qi* thresholds applied to the responses to white stimuli (Qi > 1 for all flash protocols except (c) which employed fewer stimulus repeats, where we used

*Qi* > 1.5, and for AMPA/KA puffs, where we used Qi > 3). The consistency index (Ci) is defined as the ratio between the variance of the mean and the mean of the variance across n=8 to 10 stimulus trials [33]. ROIs with *Ci* ≥ 0.2 were considered to show consistent light-evoked Ca^2+^ responses over time. For all experiments involving light stimuli, only ROIs that passed both criteria were included for further analysis.

Depending on the stimulus protocol, we determined additional parameters for each ROI: We calculated the **spectral contrast** preference, *SC = (F_Area{G)_ - F_Area{uv)_)/(F_Area{G)_ + F_Area{uv)_*), using the *F_Area_* for the responses to green and UV flashes (protocol (b)). The dark-light index, *DLi = (F_Area{B)_-F_Area{D)_)/(F_Area{B)_+F_Area{D)_*) [32], was determined using the *F_Area_* for the responses to bright and dark white flashes (protocol (c)).

The data recorded with the coloured noise stimulus (protocol (d); *cf.* Fig. 4) were analysed by calculating the negative transient-triggered average from the de-trended and z-normalized Ca^2+^ traces, weighted by the transients' amplitudes, yielding a temporal receptive field (time kernel) for each ROI. A ROI was considered light-responsive if the maximum amplitude of the kernel (*A_lrf_*) for green and/or UV was *A_lrf_*> 2 s.d. of the noise. All kernels were then normalized to 1. We then calculated the correlation between ROIs present in the same field either for the full Ca^2+^ traces or for negative events (with amplitudes < -2 s.d. of the noise) in a time window of -750 to 250 ms around the event (at 0 ms). The mean correlation for each field was then used for further analysis. An equivalent approach was used to analyse the data recorded with the binary noise (protocol (e); *cf.* Fig. 6); with ROIs considered responsive if *A_lrf_*> 3 s.d. noise. A periodogram was generated by applying a discrete Fourier transform (DFT) to the time-series of each kernel without zero padding. The power spectral densities at each frequency component followed approximately a log-normal distribution, and so to improve Gaussianity (assumed in the subsequent t-tests), a log transform was applied to each periodogram, and the transformed data was used for statistical comparisons.

### Statistics

All statistical tests (except for the ones for the periodograms) were performed using the Wilcoxon signed-rank test or the Wilcoxon rank-sum test. Alpha was set to 0.05 and p-values (*p*) < 0.05 were considered as significant (*), *p* < 0.01 (**), *p* < 0.001 (***). For multiple comparisons, Bonferroni correction was used and *p* < 0.025 was considered as significant (+), *p* < 0.005 (++), *p* < 0.0005 (+++). For periodograms, a dependent sample t-test was computed for each positive frequency component and Bonferroni correction was used (15 comparisons, *cf.* Fig. 6). Spearman rank correlation test was used to estimate the correlation between negative events and distance along the slice (*cf.* Fig. 4) as well as the relationships between *DLi, SC*, slice position and *F*_0_ (*cf.* Suppl. Information Fig. 1). Differences between dorsal and ventral *DLi* were assessed with t-test and Bartlett test. Errors are given as standard error of the mean (SEM), median absolute deviation (MAD) or standard deviation (s.d.).

### *Modelling voltage and Ca*^*2*^*+ spread across HC dendrites*

To evaluate the voltage and Ca^2+^ spread across HC dendrites (Fig. 5), we built a biophysically realistic model using the simulation language NeuronC [63]. To this end, we reconstructed a HC dendritic branch and its cone contacts from a published EM dataset (e2006; [54]) using Knossos (https://knossostool.org). The model includes AMPA-type glutamate receptors at the cone synapses and voltage-gated Ca^2+^ and K+ channels modelled as Markov state machines with different densities for tips and dendrites (for parameters, see Table 4). Photoreceptors are already pre-defined in NeuronC; they were modelled as single compartments that included voltage-gated Ca^2+^ and Ca^2+^-activated Cl^-^ channels. We adjusted channel densities such that the model's membrane voltage stays in a physiologically plausible range but did not further tune the model. The model code is available via http://eulerlab.de/data/.

**Table 4.** Parameters of biophysical model of HC dendritic branch

|  |  |  |  |
| --- | --- | --- | --- |
| Rm | $[\Omega \text{ cm}^2]$ | | 2,500 |
| Ri ($\Omega \text{ cm}^2$) | $[\Omega \text{ cm}^2]$ | | 200 |
| Channel densities |  |  |  |
| L-type $\text{Ca}^{2+}$ channels | $[\text{S}/\text{cm}^2]$ | Soma and proximal dendrites | 3e-4 |
|  |  | Distal dendrites | 1e-3 |
| $\text{K}^+$ channels | $[\text{S}/\text{cm}^2]$ | Soma and proximal dendrites | 1e-5 |
|  |  | Distal dendrites | 1e-5 |

## Supplemental Information for the manuscript titled: "Local signals in mouse horizontal cell dendrites"

### *GABAa* *receptor activation modulates the intracellular Ca*^*2*^*+ level in distal HC tips*

Horizontal cells are GABAergic [13] but the role of GABA for the HC feedback to cones is still controversial [16]. In mouse, cones do not express ionotropic GABA receptors [12], while HCs do [64]. Therefore, it has been proposed that GABA release from HCs activates GABAa autoreceptors and thereby modulates ephaptic and pH-mediated feedback to cones [13,14].

To test if GABA auto-reception affects Ca^2+^ signals in HC processes we puff-applied the GABAa receptor agonist muscimol while presenting light flashes (Suppl. Fig. 3A,B). Despite some variability, muscimol caused on average a small but significant increase in *F*0 (by 0.32 ± 0.13 s.d., p=0.011 for muscimol vs. control; n=20 ROIs from 4 slices, 2 mice; Suppl. Fig. 3C and Suppl. Table 1) which was reversible (0.03 ± 0.15 s.d., p=0.007 for muscimol vs. wash-out). The size of the light responses did not change significantly (Suppl. Fig. 3D,E, Suppl. Table 1). That Ca^2+^ levels increase upon GABAa (auto-)receptor activation is consistent with earlier reports that suggested high intracellular Cl^-^ levels in HC dendritic tips due to the expression of Na+/K+/Cl^-^ co transporters [65,66]. Thus, it is likely that GABAa receptor activation caused a Cl^-^ efflux, with the resulting depolarisation activating voltage-gated Ca^2+^ channels (VGCCs). We also puff-applied the GABAa receptor antagonist gabazine but we did not find any consistent effects (n=33 ROIs from 4 slices, 2 mice; Suppl. Fig. 3F-J, Suppl. Table 1), in line with results for mouse cone axon terminals (for discussion, see [12]). Our data provide evidence that GABA auto-reception plays a role in modulating the activity in distal HC processes and, thus, in shaping cone output [13,30].

### Contrast encoding in the dendritic tips of HCs

Mouse cones differ in their encoding of achromatic contrast depending on their location on the retina [32]: dorsal M-cones respond equally well to bright and dark flashes (positive and negative contrasts, respectively), whereas ventral S-cones prefer dark over bright contrasts. We tested if this contrast coding is preserved in HC dendrites (Suppl. Information Fig. 1) by presenting bright and dark flashes of different “colours” (Suppl. Information Fig. 1A-C, see “contrast and colour” protocol in Methods) and calculating for each ROI the spectral contrast (SC) preference and its preference for dark over bright contrasts ("dark-light index", *DLi*, Methods). The majority of HC ROIs generally responded preferentially to dark over bright stimuli (green: *ΔF_dark_* = 1.44 ± 0.60 vs. *ΔF_bright_ =* 0.62 ± 0.46, p=8.389·10^-9^, n=57; UV: *ΔF_dark_* = 1.63 ± 0.56 vs. *ΔF_bright_ =* 0.55 ± 0.35, p=2.288·10^-25^, n=99) (Suppl. Information Fig. 1A-D). Nevertheless, the *DLi* distribution for dorsal ROIs was broader than that for ventral ROIs and shifted towards zero (means: -0.51 vs -0.70, p = 0.0093, t-test; variance: 0.22 vs. 0.06, p=0.000004, Bartletts test; Suppl. Information Fig. 1E). In general, the *DLi* was independent on *SC* (Suppl. Fig. 5F) and *F_0_* (Suppl. Fig. 5G). Overall, the contrast preference in HC processes shows a trend along the dorso-ventral retina axis that is reminiscent of what was earlier shown for cones [32], arguing that distal HC dendrites inherit properties of local cones.

**Supplemental Table 1.** Pharmacology for GABAa auto-receptors on HCs. Muscimol, GABAa receptor agonist; gabazine, GABAa receptor antagonist; Ca^2+^ baseline (*F_0_*), amplitude (*ΔF*) and area-under-the-curve (*F_Area_*) of light-evoked Ca^2+^ responses. Wilcoxon signed-rank test.

|  | Number of mice/slices/ROIs | control | drug | wash |
| --- | --- | --- | --- | --- |
| muscimol puff application |  |  |  |  |
| $F_0$<br>[s.d.] | 2/4/20 | 0 | $0.323 \pm 0.126$<br>(p=0.011 +) | $0.029 \pm 0.149$<br>(p=0.007 +) |
| $\Delta F$<br>[s.d.] | | $1.0166 \pm 0.061$ | $0.891 \pm 0.102$<br>(p=0.330) | $0.762 \pm 0.087$<br>(p=0.040) |
| $F_{Area}$<br>[a.u.] | | $0.915 \pm 0.067$ | $0.707 \pm 0.110$<br>(p=0.154) | $0.649 \pm 0.086$<br>(p=0.452) |
| gabazine puff application |  |  |  |  |
| $F_0$<br>[s.d.] | 2/4/33 | 0 | $0.0622 \pm 0.076$<br>(p=0.525) | $0.024 \pm 0.060$<br>(0.437) |
| $\Delta F$<br>[s.d.] | | $0.981 \pm 0.0762$ | $0.874 \pm 0.0816$<br>(0.126) | $0.733 \pm 0.067$<br>(p=0.009 +) |
| $F_{Area}$<br>[a.u.] | | $0.822 \pm 0.079$ | $0.751 \pm 0.081$<br>(p=0.296) | $0.606 \pm 0.0679$<br>(p=0.027) |

**Supplementary Information Figure 1.**
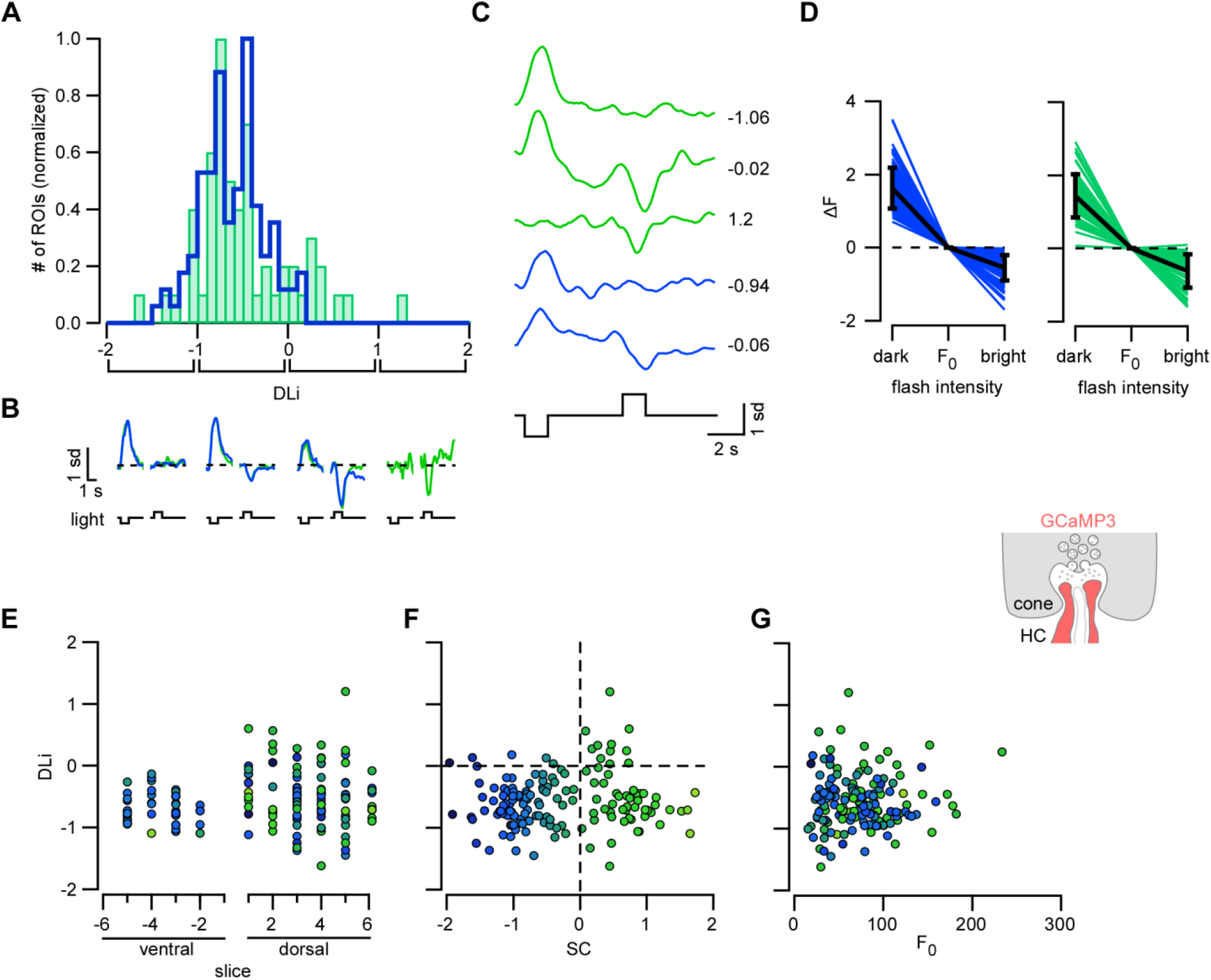
Contrast preference of Ca^2+^ responses in HC processes. **A.** Histogram of dark-light index distribution (*DLi;* see Methods) for green (SC>0; n=57 ROIs) and UV (SC<0; n=99 ROIs). **B.** Averaged Ca^2+^ signal in response to green and UV, dark and bright flashes for different *DLi* intervals (averages of n=8 trials). **C.** Exemplary green and UV ROIs responding to dark and bright flashes (averages of n=8 trials). Values indicate *DLi*. **D.** Response amplitudes (*ΔF*) for UV (left) and green ROIs (right) to dark and bright flashes (*F_0_*, baseline). **E-G.** *DLi* plotted as a function of slice position (E), *SC* (F) and baseline (*F_0_*, G). Colours reflect *SC* preference of each ROI. E. Error bars indicate s.d.

